# Systematic Development of a Compact Genome-Editing Tool Leveraging the TAM-Independent DNA Nuclease TasR in *Bacillus subtilis*

**DOI:** 10.64898/2026.07.15.738680

**Authors:** Xiaoyan Tang, Jie Gao, Hengyi Wang, Xiaomo Wei, Xirui Zhou, Xinyao Pan, Yuhao Wang, Mingxin Li, Qi Li

## Abstract

*Bacillus subtilis* is a core microbial chassis in biomanufacturing, and establishing efficient gene editing technologies is key to engineering this strain. In conventional CRISPR gene editing technologies, the large size of DNA nucleases leads to difficulties in plasmid construction, low transformation efficiency, and cumbersome multi-round editing operations; therefore, developing miniature gene editing tools can effectively address these issues. Although our group previously established a miniature gene editing tool based on IscB in *B. subtilis* SCK6, IscB relies on the 5′-CAGGAA-3′ TAM recognition sequence, and 83.36% of the genes in the SCK6 genome harbor no or only one TAM sequence, indicating a bottleneck of restricted editing for IscB in this strain. The novel miniature DNA nuclease TasR does not require a TAM sequence and can thus compensate for the limitation of IscB; however, the applicability of TasR in *B. subtilis* remains unknown. Therefore, this study first constructed a single plasmid, pBsuTasR, capable of expressing TasR and its guide RNA (tigRNA), which enabled gene deletion of regular-sized fragments in SCK6 with editing efficiencies of 21.7%– 78.3%. Subsequently, the capacity of TasR to delete a long DNA fragment (169.9 kb) was evaluated, and it was found that under the guidance of a single tigRNA, the deletion efficiency was 21.73%, whereas after optimizing to two tigRNAs, the efficiency increased to 39.13%. Furthermore, the gene integration capability of pBsuTasR was further tested, and TasR was able to integrate the *aprN* gene into the *amyE* locus at an efficiency of 13.3% under the guidance of a single tigRNA, and after increasing to two tigRNAs, the integration efficiency increased to 91.3%. In terms of iterative genome editing, this study developed the pBsu-SRP (Scissors-Rock-Paper) iterative editing system, which automatically cures the editing plasmid from the previous round while performing a new round of gene editing, with sequential gene deletion efficiencies of 4.34%–26.08%, and using this system, the editing cycle can be shortened from 4N days by the conventional method to 3N+1 days. Subsequently, the pBsu-SRP system was successfully used to achieve the integration of two and three copies of the *mCherry* fluorescent reporter gene in SCK6, and it was found that the fluorescence intensity increased with the copy number. Finally, this study also explored the escape of SCK6 from TasR cleavage and found that mutations in the tigRNA sequence are the cause of the escape. In summary, this study constructed a novel miniature genome editing system in *B. subtilis* using the TAM-independent nuclease TasR as the core component. This system can not only provide an efficient technical tool for genetic manipulation of industrial microorganisms, but also offer new instrumental support for the iterative engineering and functional optimization of chassis cells in biomanufacturing.

## 1. INTRODUCTION

The global biomanufacturing industry is accelerating its transition toward green and low-carbon operations, with synthetic biology emerging as a core driving force for upgrading traditional industrial processes^[1]^.*Bacillus subtilis*, a generally recognized as safe (GRAS) Gram-positive industrial microorganism, has been extensively employed in the large-scale production of enzyme preparations, bulk chemicals, and natural products, owing to its robust protein secretion capacity, well-characterized genetic background, and favorable fermentation properties^[2,3]^. High-efficiency and precise genome editing tools constitute the foundation for constructing high-performance industrial strains^[4,5]^, directly determining the efficiency and duration of metabolic engineering modifications^[6–8]^.

The advent of RNA-guided DNA nuclease technologies has revolutionized the field of microbial genome editing^[9–11]^. Among these, the CRISPR-Cas9 and CRISPR-Cpf1 systems have been widely applied in *B. subtilis*, enabling the establishment of both single-plasmid and dual-plasmid editing platforms^[12–14]^. However, conventional CRISPR systems still exhibit notable limitations. First, the large molecular sizes of Cas9 and Cpf1 (1000–1400 aa) impede efficient loading onto single-plasmid vectors, leading to low transformation efficiencies. Although dual-plasmid systems mitigate the cargo burden, they introduce greater operational complexity and necessitate additional antibiotic selection markers. Second, for large-fragment DNA deletions, CRISPR-Cas9 generally requires two guide RNAs (gRNAs) to achieve deletions at the 100-kb scale, whereas CRISPR-Cpf1 can mediate large-fragment deletions with a single gRNA; however, this requires prior chromosomal integration of the Cpf1-encoding gene, resulting in a protracted and laborious preparatory workflow. Moreover, iterative multi-round editing is constrained by common bottlenecks, including a limited repertoire of selectable markers and cumbersome plasmid curing procedures^[15–18]^. Consequently, the development of miniature nuclease-based genome editing tools with reduced molecular footprints and streamlined workflows has emerged as a critical direction for overcoming the inherent limitations of conventional CRISPR systems.

In recent years, a series of smaller RNA-guided DNA nucleases have been successively discovered. In 2021, IscB (∼500 aa) was first demonstrated as an evolutionary ancestor of CRISPR-Cas9 possessing programmable double-stranded DNA cleavage activity; with a molecular weight only one-third that of Cas9, it substantially alleviates the vector cargo burden^[19]^. To date, IscB has enabled efficient gene editing and base correction in human cells and murine disease models, and preliminary applications have also been reported in a limited number of Gram-negative strains, including *Escherichia coli* MG1655^[20,21]^. Our group has previously established IscB-based editing systems in *B. subtilis* SCK6 using both wild-type IscB and an enhanced variant, enIscB; however, IscB was found to strictly require the 5′-CAGGAA-3′ target-adjacent motif (TAM), rendering more than half of the *B. subtilis* genes inaccessible to effective targeting. Furthermore, its host adaptability remains considerably limited, falling short of the requirements for genome-scale metabolic engineering^[15]^. Accordingly, the development of novel miniature nuclease editing tools with sequence-unrestricted targetability is of pressing practical necessity for overcoming current technological bottlenecks and expanding the genetic operability boundaries of *B. subtilis*.

TasR, the core nuclease of the TIGR-Tas system reported in 2024, provides a new approach to address the above limitations^[10]^. TasR is the effector nuclease of the TIGR system and belongs to a class of miniature RNA-guided DNA endonucleases. It consists of only ∼350 amino acids (∼40 kDa) and harbors an N-terminal RuvC nuclease domain and a C-terminal conserved Nop domain. TasR forms a homodimer through a coiled-coil domain and assembles with a 36-nt tigRNA into a functional complex. It does not require a TAM motif; instead, it targets double-stranded DNA through the dual-spacer sequence of the tigRNA, generating 8-nt 3′ overhangs. TasR is not only substantially smaller than the conventional Cas9 protein, but also more compact than the miniature nuclease IscB (∼500 aa, ∼55 kDa) previously used by our group, offering advantages in plasmid loading and delivery, and can be reprogrammed to achieve precise DNA cleavage^[10,22]^.

In contrast to the TAM-dependent targeting bottleneck of IscB, this motif-independent targeting property of TasR theoretically enables coverage of any genomic locus, fundamentally addressing the core limitation of IscB’s restricted editable space. To date, studies on TasR have been predominantly focused on gene editing applications in eukaryotic systems, and no systematic characterization of its editing properties or applications in *B. subtilis* SCK6 has been reported^[10,23,24].^

In summary, this study systematically investigates the gene editing performance and application potential of TasR in *B. subtilis* SCK6. First, a single-plasmid editing system, pBsuTasR, was constructed to assess the toxicity of TasR in *B. subtilis* SCK6. Subsequently, the single-gene deletion capability of TasR was evaluated, followed by verification of its efficiency in deleting long DNA fragments. The integration efficiency of exogenous genes mediated by TasR was then tested. On this basis, an iterative genome editing system based on the “Scissors-Rock-Paper (SRP)” strategy was established, enabling multi-round continuous gene editing without counter-selection or serial passage-dependent plasmid curing. The feasibility of multi-copy integration was further validated using the pBsu-SRP system with an *mCherry* fluorescent reporter. Finally, the patterns and mechanisms underlying *B. subtilis* escape from TasR cleavage were preliminarily explored. This work expands the repertoire of miniature nuclease-based genome editing tools in *B. subtilis*, provides novel technical support for efficient genetic engineering of industrial microorganisms, and holds significant implications for advancing the high-quality development of the green biomanufacturing industry.

## 2 Materials and Methods

### 2.1 Strain cultivation

The strains used in this study are listed in Supplementary S1. *Escherichia coli* DH5α was used as the cloning host for plasmid construction. The constructed strains were *E*. *coli* MG1655 and *Bacillus subtilis* SCK6. Strains were cultured in LB medium (0.5% [w/v] yeast extract, 1% [w/v] tryptone, 1% [w/v] NaCl) at 37 ℃ and 220 rpm. For plasmid selection, kanamycin was added to LB medium at 50 μg/mL for E. coli MG1655 and 20 μg/mL for *B. subtilis* SCK6. *B. subtilis* SCK6 competent cells were prepared in YN medium (0.7% [w/v] yeast extract, 1% [w/v] tryptophan, 0.5% [w/v] NaCl, 0.3% [w/v] beef extract) supplemented with 1.5% xylose. Strains were stored in 20% glycerol at -80 ℃. For revival, strains were streaked on LB agar plates. Single colonies were inoculated into liquid LB medium and cultured overnight at 37 ℃ and 220 rpm for 12 h.

### 2.2 Reagents and Enzymes

Restriction enzymes used in this study were purchased from Thermo Fisher Scientific (Waltham, MA, USA). DNA amplification was performed using DNA polymerase KOD plus Neo (Toyobo Co., Ltd., Osaka, Japan) and DNA polymerase 2× Phanta Flash Master Mix (Vazyme Biotechnology Co., Ltd., Nanjing, China). 2× Es Taq Master Mix (Dye) (Jiangsu Cowin Biotech Co., Ltd., Taizhou, China) was used for colony polymerase chain reaction (PCR) of *E. coli* MG1655, and 2× Rapid Taq Master Mix (Dye) (Vazyme Biotechnology Co., Ltd., Nanjing, China) was used for colony PCR of *B. subtilis* SCK6. Plasmid extraction and DNA purification kits were purchased from Tiangen Biotech Co., Ltd. (Beijing, China) and used according to the manufacturer’s instructions. Plasmid construction was conducted via the ClonExpress One-Step Cloning Kit (catalog numbers C116 and C112, Vazyme Biotech Co., Ltd., Nanjing, China).

### 2.3 Plasmid Construction

All plasmids and primers used in this study are listed in Supplementary S1 and S2, and *E. coli* DH5α was used for the maintenance and construction of plasmid. For the construction of pBsuTasR-targetX (targetX denotes target gene, Molecular Cloud ID: MC_0102155), the vector backbone was amplified with pBsuIscB-ωRNA-targetX as the template, the coding sequences of IscB and ωRNA were replaced with sequences expressing TasR protein and tigRNA, where the TasR gene was controlled by the xylose-inducible promoter P_xylA_ and tigRNA was driven by the constitutive promoter Pveg; all obtained fragments were recombined with double-stranded DNA repair templates containing 1 kb upstream and 1 kb downstream homologous arms amplified from the *B. subtilis* SCK6 genome, and the recombination products were transformed and spread on kanamycin-resistant plates for positive strain screening, followed by colony PCR verification of recombinant plasmids, plasmid extraction and sequencing to confirm successful assembly. To construct pEc-TasR (Molecular Cloud ID: MC_0102151) and pEc-tigRNA-targetX, the plasmid backbone was amplified from pEcTnpB-2.0, and the TnpB coding sequence was substituted by the expression cassette of TasR together with the L-arabinose-inducible λ RED promoter amplified from pSC101 Cas9; the recombinant products were chemically transformed competent cells and cultured on chloramphenicol-resistant plates for screening, and transformants carrying intact TasR expression elements were identified via colony PCR, indicating complete replacement of the original TnpB gene and its promoter to generate pEc-TasR. The cleavage plasmid pEc-tigRNA-targetX (targetX denotes target gene, Molecular Cloud ID: MC_0102156) was constructed by two modifications of pTargetF: the N20 fragment of pTargetF was replaced with a 36 nt sequence (5′-AGCCA-spacerA-UGAAACCCA-spacerB-UGCG-3′), and the intrinsic gRNA scaffold sequence was simultaneously removed.

### 2.4 Preparation of competent cells and plasmid transformation

Take 10 μL of wild-type *E*. *coli* MG1655 bacterial stock stored at -80 ℃ and streak it onto an LB agar plate without antibiotics for revival. Subsequently, inoculate a single colony from the plate into fresh liquid LB medium and culture overnight at 37 ℃ with shaking at 220 rpm for 12 h. The next day, transfer 50 μL of the overnight culture into fresh liquid LB medium (a ten-fold scale-up in inoculum volume compared with the overnight culture inoculation), incubate at 37 ℃ and 220 rpm for 2 h, and then prepare competent cells in a final volume of 100 μL by a cell washing procedure. Mix the prepared plasmids with the competent cells and perform electroporation at 2.5 kV in a chilled 2 mm Gene Pulser cuvette (Bio-Rad, Hercules, USA). Immediately after electroporation, resuspend the mixture in 900 μL of fresh liquid LB medium and incubate at 37 ℃ with shaking at 220 rpm for 1 h. Subsequently, plate the incubated cells onto LB agar plates containing the appropriate antibiotic for plasmid selection and incubate at 37 ℃ overnight for 12-15 h. Calculate colony-forming units (CFU) based on the colonies grown on the plates and determine the number of plasmid transformants.

Streak the *B. subtilis* SCK6 stock stored at -80 ℃ onto an LB agar plate for revival. Subsequently, inoculate a single colony from the plate into fresh LB medium. Incubate the culture at 37 ℃ with shaking at 220 rpm for 12 h. Then, transfer the overnight culture into fresh YN medium supplemented with 1.5% xylose (initial OD600 of 1.0) and incubate at 37 ℃ and 220 rpm for 2 h to generate competent cells. Mix the prepared plasmids with 500 μL of competent cells and incubate at 37 ℃ and 220 rpm for 90 min. Subsequently, plate the incubated cells onto LB agar plates containing the appropriate antibiotic for plasmid selection and incubate at 37 ℃ overnight for 12-15 h. Calculate colony-forming units (CFU) based on the colonies grown on the plates and determine the number of plasmid transformants.

#### 2.5.1 Distribution analysis of TAM sequences for IscB in *B.subtilis*

In this study, the whole-genome annotation file (GenBank format) of the *B. subtilis* 168 strain was parsed using the Biopython toolkit. A Python script was employed to iterate through all CDS coding region features, and target genes containing or lacking the “CAGGAA” nucleotide sequence were identified by a string-matching algorithm. Gene names, start/end positions, and nucleic acid sequences were structurally stored as an Excel file using the pandas library. Meanwhile, gene coordinates were converted to the kilobase (kb) scale with the matplotlib data visualization tool, and a linear distribution map of the target genes across the whole genome was generated, thereby revealing the spatial distribution patterns of the genes corresponding to this sequence on the chromosome.

#### 2.5.2 Gene deletion and insertion analysis in B.subtilis SCK6

Add 10 µL of pBsuTasR-targetX plasmid at a known concentration to 500 μL of *B. subtilis* competent cells, mix thoroughly, and incubate at 37 ℃ and 220 rpm for 90 min. The mixture was then spread onto LB agar plates containing 3% xylose and kanamycin (20 μg/mL) and incubated overnight at 37 ℃. Twenty-three transformants were randomly selected from the selection plates and verified by PCR and DNA sequencing, with the wild-type strain serving as the control. The deletion primers were complementary to the sequences ∼50 bp upstream and ∼50 bp downstream of the genomic homology arms (∼1 kb). Successfully edited strains were stored at -80℃ at a final concentration of 20% glycerol.

#### 2.5.3 Design and construction of the Rock-Paper-Scissors gene editing system

The “Rock-Paper-Scissors” strategy required three plasmids, namely pBsu-Scissors (Scissors), pBsu-Rock (Rock), and pBsu-Paper (Paper), and the system composed of these three plasmids was designated the pBsu-SRP system^[25,26]^. Each plasmid contained basic elements including a replicon, a gene expressing TasR, homology arms for gene editing, and a selection marker gene. The selection markers used in *B. subtilis* were the chloramphenicol resistance gene *CmR*, the kanamycin resistance gene *KanR*, and the tetracycline resistance gene *TcR*. Notably, each plasmid carried two tigRNAs: tigRNA1 targeted the gene of interest, and tigRNA2 targeted the resistance gene on the plasmid from the previous round of editing. In addition, three identical resistance gene editing sites were designed on the previous-round editing plasmid, located on both sides of the resistance gene and within the resistance gene sequence. The detailed construction methods for each plasmid are as follows:

Scissors plasmid pBsu-Scissors (Molecular Cloud ID: MC_0102154): According to the method described in section 2.3, the N18 sequence of tigRNA1 was replaced with the target gene, and the upstream and downstream homology arms of this gene were added to construct the pBsuTasR-targetX plasmid. Then, using pBsuTasR-targetX as the backbone, tigRNA2 and the chloramphenicol resistance gene *CmR* (derived from plasmid pHT-XCR6; this resistance marker was used for plasmid selection in *B. subtilis*) were inserted between the *B. subtilis* replicon rep pE194ts and the homology arms. In addition, the ampicillin resistance gene *AmpR* (derived from plasmid pHT-XCR6; this resistance marker was used for plasmid selection in *E. coli*) was added between the homology arms and the *E. coli* replicon p15A.

Rock plasmid pBsu-Rock (Molecular Cloud ID: MC_0102153): According to the method described in section 2.3, the N18 sequence of tigRNA1 was replaced with the target gene, and the upstream and downstream homology arms of this gene were added to construct the pBsuTasR-targetX plasmid. Then, using pBsuTasR-targetX as the backbone, tigRNA2 was inserted between the *B. subtilis* replicon rep pE194ts and the homology arms. This plasmid retained the kanamycin resistance marker for selection in both *E. coli* and *B. subtilis*.

Paper plasmid pBsu-Paper (Molecular Cloud ID: MC_0102152): According to the method described in section 2.3, the N18 sequence of tigRNA1 was replaced with the target gene, and the upstream and downstream homology arms of this gene were added to construct the pBsuTasR-targetX plasmid. Then, using pBsuTasR-targetX as the backbone, tigRNA2 and the tetracycline resistance gene *TcR* (derived from plasmid phy300plk; this resistance marker was used for plasmid selection in *B. subtilis*) were inserted between the *B. subtilis* replicon rep pE194ts and the homology arms. In addition, the ampicillin resistance gene *ampR* (derived from plasmid pHT-XCR6; this resistance marker was used for plasmid selection in *E. coli*) was added between the homology arms and the *E. coli* replicon p15A.

After the three plasmids of the complete system were constructed, when changing the gene to be edited, it was only necessary to replace the N18 sequence of tigRNA1 with the target gene sequence and to replace the homology arms with the upstream and downstream homology arms of the target gene.

#### 2.5.4 Plasmid curing

To verify the feasibility of the pBsu-SRP system plasmids, after each round of gene editing, antibiotic selection was carried out to obtain knockout strains harboring only the editing plasmid for the next round. To cure the editing plasmid pBsu-Scissors-targetA, a colony of the edited clone containing pBsu-Scissors-targetA was inoculated into 5 mL of LB medium supplemented with kanamycin (20 μg/mL) and chloramphenicol (25 μg/mL). The culture was incubated overnight with shaking at 220 rpm. The next day, 10 μL of the culture was transferred into LB medium containing kanamycin (20 μg/mL) and incubated at 50 ℃ and 200 rpm for 8 h, and this passage was repeated for three consecutive times. Subsequently, approximately 10 μL of the cell suspension was diluted and spread onto LB plates containing both kanamycin (20 μg/mL) and chloramphenicol (25 μg/mL), as well as onto LB plates containing kanamycin (20 μg/mL) only, and incubated overnight at 37 ℃. Colonies that grew on the LB plates containing only kanamycin were randomly picked and patched onto LB plates containing kanamycin (20 μg/mL) plus chloramphenicol (25 μg/mL) and onto LB plates containing only kanamycin (20 μg/mL) for screening. Colonies sensitive to chloramphenicol had lost pBsu-Scissors-targetA. The same method could be used for curing pBsu-Rock-targetB and pBsu-Paper-targetC.

#### 2.5.5 Phenotypic verification of *spo0A*, *amyE*, *sacB*, and *aprE*knockout strains

The control strain, wild-type *B. subtilis* SCK6, and the experimental strain, *B. subtilis* SCK6 Δ*spo0A*, were taken from glycerol stocks, and 10 μL of each bacterial suspension was spread onto LB agar plates for revival. Single colonies were picked and inoculated into liquid LB medium for overnight culture. An LB agar plate was evenly divided into two halves; 10 μL of fresh culture from each strain was taken, diluted to 10^-4^, and spread onto the two halves of the plate, respectively. After overnight incubation, colony morphology was observed. Similar methods were used for phenotypic verification of *B. subtilis* SCK6 Δ*amyE*, *B. subtilis* SCK6 Δ*sacB*, and *B. subtilis* SCK6 Δ*aprE*.

#### 2.5.6 Fluorescence Intensity Measurement

To eliminate interference from plasmid-encoded *mCherry* expression, plasmids were cured from the chromosomally integrated strains, and the genotypes of the strains were confirmed by PCR. After plasmid curing, the *mCherry*-integrated *B*. *subtilis* strain was cultured overnight in LB medium, and 1% volume of the overnight culture was inoculated into 50 mL fresh LB medium. Culture samples were harvested at 4 h, 6 h, 8 h, 10 h, 12 h, 24 h and 48 h to determine OD₆₀₀ values and fluorescence intensity. The fluorescence signal of *mCherry* was detected with an excitation wavelength of 587 nm and an emission wavelength of 610 nm. In parallel, cell pellets collected by centrifugation were visually inspected for *mCherry* fluorescence. Wild-type *B. subtilis* SCK6 was used as the blank control.

#### 2.5.7 Analysis of Escape Patterns of *B*. *s*ubtilisagainst TasR-Mediated Cleavage

Following the procedures described in section 2.3, the fragment without donor DNA was amplified using plasmid pBsuTasR-*amyE*-tigRNA as the template to serve as the vector backbone. This backbone was recombined and assembled with the newly designed second tigRNA to construct plasmid pBsuTasR-*amyE*-tigRNA2. The successfully constructed pBsuTasR-*amyE*-tigRNA2 plasmid was transformed into prepared competent cells of *B. subtilis*. After 1.5 h of recovery culture, the cell mixture was spread onto LB agar plates supplemented with kanamycin and 3% xylose, followed by overnight incubation until distinct single colonies formed. Single colonies were randomly picked, and the tigRNA1-TasR fragment as well as the tigRNA2 fragment were amplified separately via PCR. The purified PCR products were subjected to Sanger sequencing, and the sequencing data were aligned with the plasmid map to summarize and analyze the patterns through which *B. subtilis* escapes TasR-dependent cleavage.

#### 2.6.1 Evaluation of gene deletion efficiency of TasR in *E*. *coli*MG1655

Since the TasR plasmid carries the promoter of the λ-Red system, 10 mM arabinose was added to the culture of *E*. *coli* harboring the pEc-TasR plasmid during cultivation to the logarithmic phase to induce expression of the λ-Red system. Subsequently, 100 μL of the prepared competent cells were mixed with 100 ng of the tigRNA-targetX plasmid and 400 ng of the tigRNA-targetX-HR plasmid^[21]^. After electroporation and incubation, the cells were spread onto LB agar plates containing chloramphenicol and spectinomycin and incubated overnight at 37 ℃ in a constant-temperature incubator. The next day, 15 transformants were randomly selected for colony PCR verification using primers complementary to the sequences ∼50 bp upstream and downstream of the genomic homology arms. The PCR products were sequenced to confirm successful gene deletion, and the corresponding wild-type strain was used as a control.

#### 2.6.2 Workflow for verifying targeted gene escape from cleavage in *E. coli* MG1655

The procedure of the escape rate assay was as follows: 100 µL of competent cells of *E. coli* MG1655 carrying pEc-TasR were mixed with the ptigRNA-TasR-TargetX series plasmids. After electroporation, the mixture was suspended in 900 µL of fresh LB broth and incubated at 37 ℃ for 1 h. The resulting mixture was plated onto solid medium and incubated overnight at 37 ℃. The colonies were counted to calculate the CFU of the experimental group, and *E. coli* MG1655 containing the ptigRNA-null plasmid (does not target any site on the genome) was regarded as the control group. (Escape rate = CFU of the ptigRNA-TasR-TargetX series plasmids in *E. coli* MG1655 containing TasR / CFU of the ptigRNA-null plasmid in *E. coli* MG1655 containing TasR).

### 2.7 Statistics and reproducibility

All the statistical data were calculated by using the GraphPad Prism 10.1.2 and all experiments were performed at least three times (n = 3 independent experiments). Data are shown as mean ± standard error mean (SEM). The statistical significance of differences between the two groups was determined by using one-way ANOVA analysis.

## 3. Results and Analysis

### 3.1 The DNA nuclease TasR exhibits broader recognition sites in *B.subtilis*SCK6

*B. subtilis* SCK6, a food-grade industrial microbial chassis, has been widely employed in enzyme preparations, bulk chemicals, and natural products biosynthesis^[1–3]^. The development of compact gene editing tools is of great significance for its metabolic engineering: compared with conventional CRISPR-Cas9 (∼1,368 aa), miniature nucleases can significantly reduce the plasmid burden, improve the construction efficiency and transformation success rate of multiplex gene editing vectors, and are better suited to the genetic manipulation characteristics of *B. subtilis*^[15–17]^.

Our group has previously established a compact gene editing system based on the DNA nuclease IscB (496 aa) in *B. subtilis* SCK6. IscB mediates DNA double-strand breaks (DSBs) by forming a ribonucleoprotein complex with ωRNA, and its cleavage activity is strictly dependent on the 5’-CAGGAA-3’ target-adjacent motif (TAM). Although IscB offers the advantage of its compact size, its strict dependence on the TAM sequence constitutes an inherent bottleneck that limits the editable genomic range. We analyzed the TAM sequences in *B. subtilis* SCK6, and the results revealed that: The genome contains 4237 annotated genes, of which 2351 (55.49%) lack IscB TAM sequences and are therefore completely refractory to IscB-mediated targeting, while 1181 genes (27.87%) harbor only a single IscB TAM sequence, severely restricting the flexibility of editing target site selection (Fig. 1A,B,C). This constraint restricts IscB-mediated targeting to fewer than half of the functional genes in *B. subtilis*, rendering it insufficient for genome-scale metabolic engineering applications.

**Figure 1.**
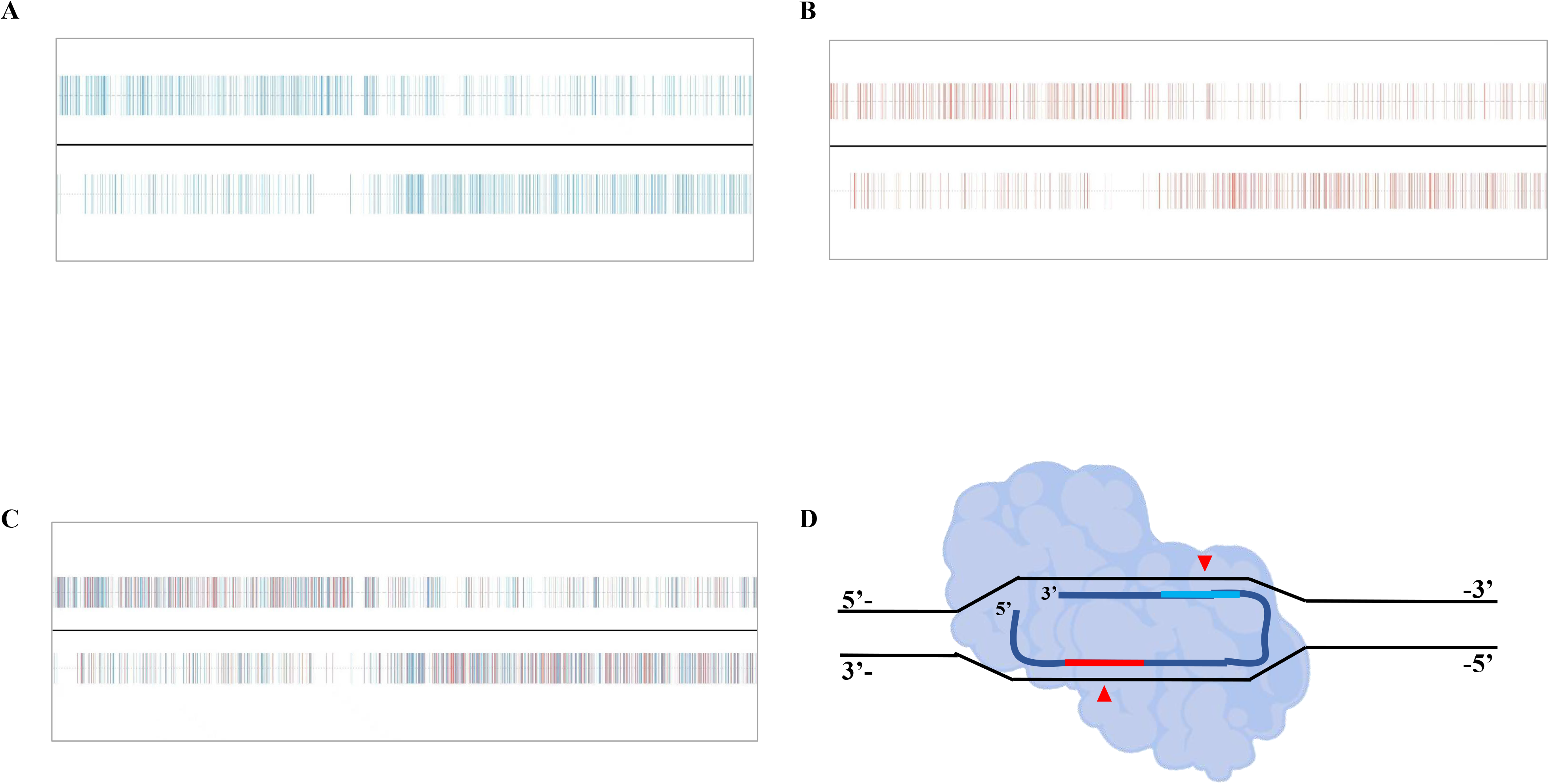
Distribution of TAM sequences for the miniature DNA nuclease IscB in *Bacillus subtilis* SCK6 and the gene editing principle of a novel miniature DNA nuclease TasR in this strain. (A) Distribution of genes lacking a TAM sequence in the SCK6 genome. (B) Distribution of genes containing a single TAM sequence in the SCK6 genome. (C) Distribution of genes containing either a single TAM sequence or no TAM sequence in the SCK6 genome. (D) Schematic diagram of the gene editing principle of TasR in SCK6.

To overcome the TAM-dependent bottleneck of IscB and further develop compact gene editing tools with higher target selectivity, this study introduces TasR, a novel TAM/PAM-independent DNA nuclease. TasR requires only a 36-nt tigRNA to form a ribonucleoprotein complex, enabling specific recognition and cleavage through complementary base pairing between the guide sequence and target DNA, independent of TAM/PAM sequences (Fig. 1D). This property enables TasR to target any gene in the *B. subtilis* SCK6 genome, covering genes that are inaccessible to IscB, while simultaneously providing more target options for genes harboring only a single TAM sequence. Therefore, the development of a novel TasR-based gene editing tool for *B. subtilis* SCK6 is expected to overcome the limitations of IscB and provide a core tool for efficient genetic engineering of this chassis.

### 3.2 Establishment of a compact genome editing tool based on TasR in *B.subtilis*

To verify the feasibility and genome editing efficiency of TasR in *B. subtilis* SCK6 and to overcome the TAM sequence dependency of the IscB system, three non-essential genes— *spo0A*, *sacB*, and *amyE*—were selected for deletion analysis, among which *amyE* lacks a TAM sequence. The upstream and downstream homologous arm sequences (∼1 kb) of each target gene were cloned into the pBsuTasR-tigRNAX vector, yielding the genome editing plasmids pBsuTasR-*spo0A*, pBsuTasR-*sacB*, and pBsuTasR-*amyE*. Subsequently, these plasmids were individually chemically transformed into competent *B. subtilis* SCK6 cells, and 23 transformants were randomly picked to assess the gene deletion efficiency (Fig. 2A). Colony PCR screening results indicated that all three target genes yielded the expected positive bands when amplified with primers designed outside the homologous arms, as confirmed by agarose gel electrophoresis. The knock-out efficiency of each gene was 21.7%(*spo0A*), 30.4%(*sacB*) and 78.3%(*amyE*), respectively (Fig. 2B). To further confirm the gene deletions in the mutant strains, DNA sequencing was performed. The sequencing results were fully consistent with the expected sequences following deletion of the target genes. Therefore, based on the sequencing data, it was confirmed that the target genes of Δ*spo0A*, Δ*sacB* and Δ*amyE* strains had been precisely deleted, and the length of deleted fragments was 804 bp, 1422 bp and 2066 bp, respectively (Fig. 2C).

**Figure 2.**
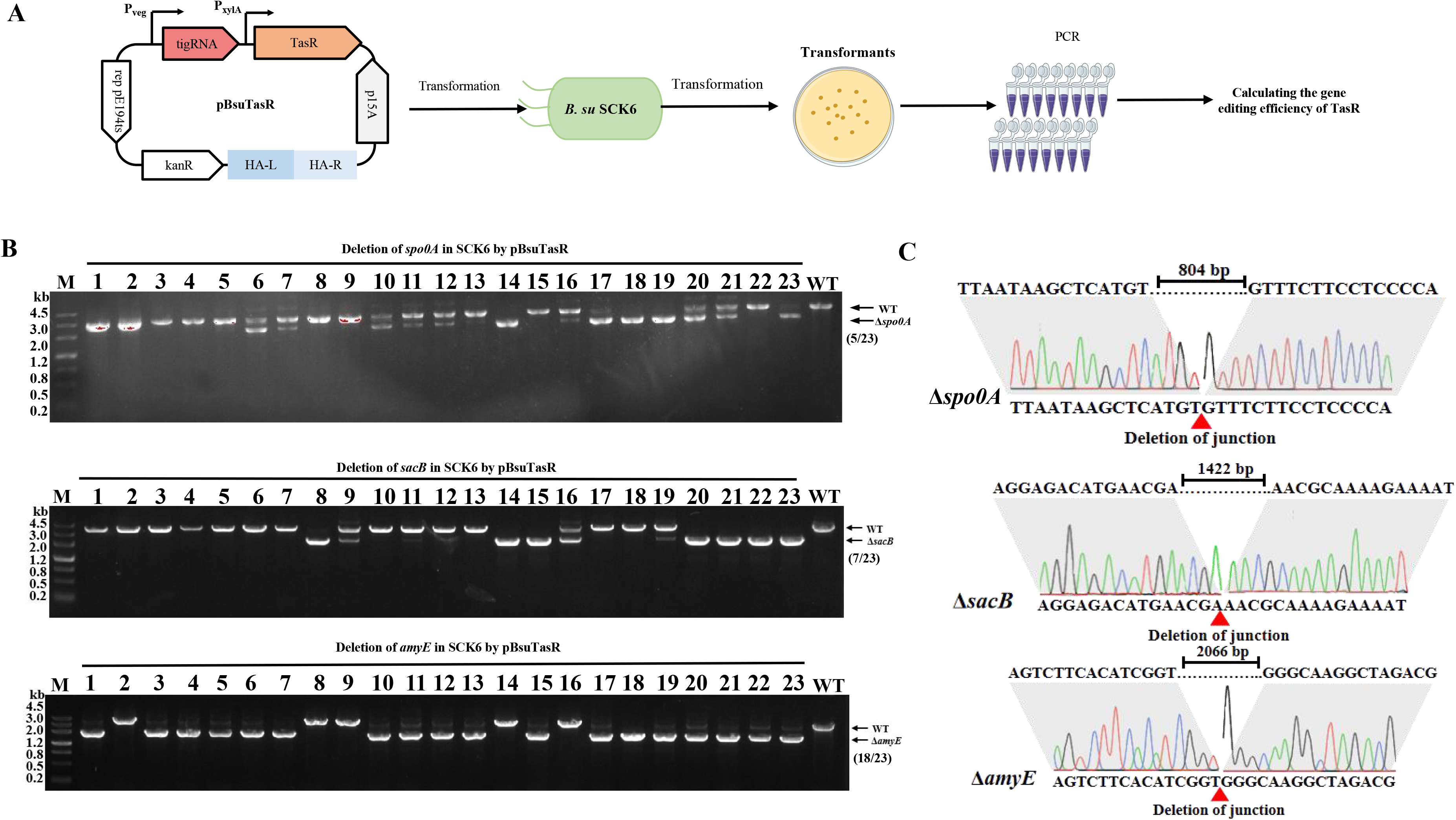
Establishment of a TasR-based genome editing system, pBsuTasR, in *Bacillus subtilis* SCK6. (A) Schematic diagram of gene deletion in SCK6 using the pBsuTasR system. (B) DNA gel images for PCR verification of *spo0A, sacB*, and *amyE* gene deletions by the pBsuTasR system. (C) DNA sequencing results confirming *spo0A*, *sacB*, and *amyE* gene deletions by the pBsuTasR system. Deletion sizes: Δ*spo0A*, 804 bp; Δ*sacB*, 1422 bp; and Δ*amyE*, 2066 bp.

In summary, the TasR-based pBsuTasR system was successfully established in *B. subtilis* SCK6, and it can accurately delete target genes with different lengths without depending on TAM sequences, with an editing efficiency of 21.7%-78.3%. Compared with the previously reported editing system of *B. subtilis* IscB, TasR also knocks out *spo0A*, and the editing efficiency of TasR is higher than that of enIscB(13.3%), and it can also efficiently edit *amyE* gene without TAM sequence, which IscB cannot achieve. These findings further substantiate that TasR completely overcomes TAM sequence dependency, enabling effective editing of genes lacking TAM sequences in *B. subtilis* SCK6. The system confers a significant advantage in target selection flexibility and provides a new technical alternative for genome engineering of this strain.

### 3.3 TasR enables deletion of long DNA fragments in *B.subtilis*

Having confirmed that TasR can accomplish conventional single-gene deletions of *spo0A*, *sacB*, and *amyE* in *B. subtilis* SCK6, we next turned our attention to the more challenging task of long DNA fragment deletions, which represents a key bottleneck in microbial editing tool optimization. In industrial chassis strain engineering, deletion of long DNA fragments spanning tens to hundreds of kilobases is frequently required to reduce metabolic burden, streamline redundant gene networks, and block competing metabolic pathways^[26,27]^. However, conventional CRISPR-based editing tools generally suffer from cumbersome workflows and low efficiency in long DNA fragment deletions, limiting the advancement of genome streamlining efforts^[15–17]^. Therefore, we selected a non-essential genomic region of 169.9 kb from *B. subtilis* SCK6 as the target, and systematically verified the deletion efficiency of long DNA fragments by TasR.

We first employed a single tigRNA targeting strategy for preliminary validation. A single recognition site was designed within the 169.9 kb region, and the ∼1 kb upstream and downstream homologous arms together with a single tigRNA element targeting this region were assembled into the pBsuTasR vector, yielding the large-fragment deletion editing plasmid pBsuTasR-169-1 (Fig. 3A). After the plasmid was chemically transformed into *B. subtilis* SCK6, 23 transformants were randomly selected for PCR and sequencing verification. The results showed that the positive deletion strain could amplify the expected size fragment, while the wild-type strain had no corresponding band because the fragment was too large, which was in line with the expectation.In efficiency, the deletion efficiency of 169.9 kb fragment mediated by tigRNA was 21.7%. DNA sequencing further confirmed that the 169.9 kb region was accurately deleted (Fig. 3B).

**Figure 3.**
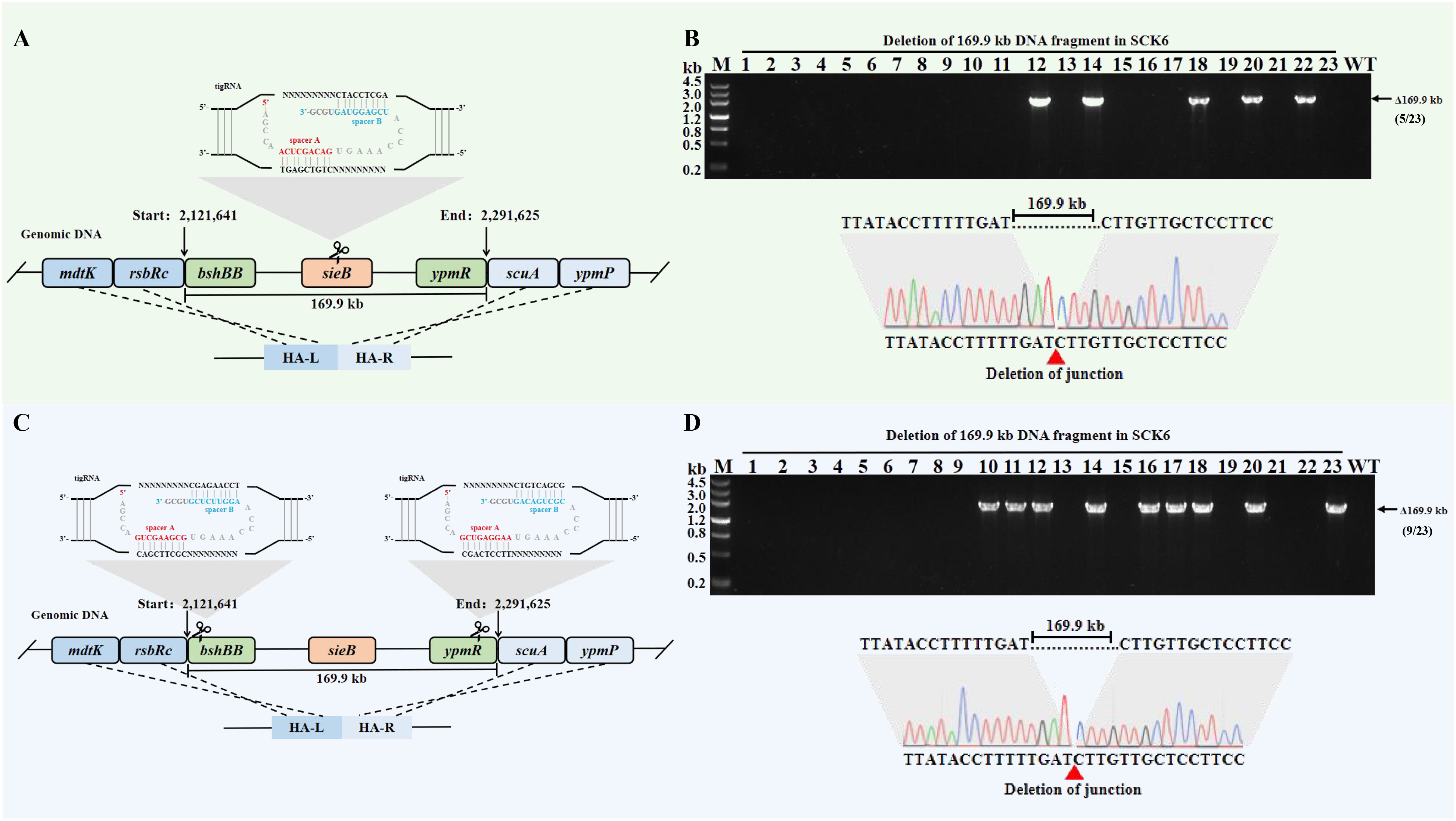
The pBsuTasR system enables deletion of large DNA fragments in *Bacillus subtilis* SCK6. (A) Schematic diagram of the single target site and homology arms for a 169.9 kb genomic fragment. (B) DNA gel image and sequencing results confirming the deletion of the 169.9 kb genomic fragment targeted by a single tigRNA. (C) Schematic diagram of the dual target sites and homology arms for the 169.9 kb genomic fragment. (D) DNA gel image and sequencing results confirming the deletion of the 169.9 kb genomic fragment targeted by dual tigRNAs.

To further improve large-fragment deletion efficiency, we posited that a “dual-nick” strategy could serve as a feasible approach to enhance gene deletion efficiency. Accordingly, we developed an optimized dual-tigRNA targeting strategy in which two cutting sites were designed at both ends of the region, separated by 169.9 kb, enabling simultaneous dual-tigRNA cleavage to increase double-strand break (DSB) efficiency. The homologous arm architecture remained consistent with that of the single-plasmid system (Fig. 3C). Using the same transformation and screening process as the single tigRNA targeting strategy, 23 transformants were randomly selected for PCR and sequencing verification. Statistics show that the deletion efficiency mediated by double tigRNA is 39.1%, and the DNA sequencing results also confirm the successful deletion of 169.9 kb genome fragment (Fig. 3D). Compared with the single-tigRNA strategy, the dual-tigRNA targeting strategy exhibited higher efficiency in long DNA fragment deletion.

In summary, TasR can delete the 169.9 kb DNA fragment in *B. subtilis* SCK6, and the double tigRNA strategy can further improve the editing efficiency. Compared with the pBsuenIscB system developed by our team in the early stage, the deletion efficiency of TasR for the same segment (39.1%) is basically the same as that of enIscB(40%)^[15]^. Compared with traditional CRISPR tools, TasR has more outstanding advantages, which are as follows: Conventional CRISPR-Cas9 typically requires a dual-plasmid system to achieve deletions exceeding 100 kb, involving multiple rounds of plasmid construction and transformation^[28]^. Although CRISPR-Cpf1 can accomplish large-fragment deletions via a single gRNA, it necessitates prior integration of the *cpf1* gene into the host genome, resulting in a lengthy preparatory modification period^[29]^. In contrast, TasR integrates the tigRNA and homologous repair elements into a single plasmid, which can be introduced into host cells via chemical transformation, substantially streamlining the workflow. More importantly, TasR is free of any PAM/TAM sequence dependency, allowing optimal target sites to be freely selected at both ends of long DNA fragments, thereby completely overcoming the target site selection limitations of conventional tools. Overall, TasR balances editing efficiency with operational simplicity, representing a promising alternative tool for genome streamlining in *B. subtilis*.

### 3.4 TasR-mediated gene integration in *B.subtilis*

To investigate the capability of TasR for site-specific integration of exogenous genes in *B. subtilis* SCK6, the *amyE* locus was selected as the integration target, and the *aprN* gene was chosen as the cargo gene. The *aprN* gene, driven by the Pveg promoter, was inserted between the upstream and downstream homologous arms of the pBsuTasR-*amyE* plasmid, yielding the integration plasmid pBsuTasR-*amyE*-*aprN* (Fig. 4A). This plasmid was transformed into *B. subtilis* SCK6, followed by overnight incubation and colony PCR verification of integration events. The results showed that TasR could successfully integrate *aprN* gene into *amyE* locus, and the integration efficiency was 56.52% (Fig. 4B), but all the integrated strains were accompanied by wild-type bands. These results demonstrate that TasR successfully integrated the *aprN* gene into the *amyE* locus; however, mixed populations were observed, indicating that editing efficiency must be further improved to eliminate this issue.

**Figure 4.**
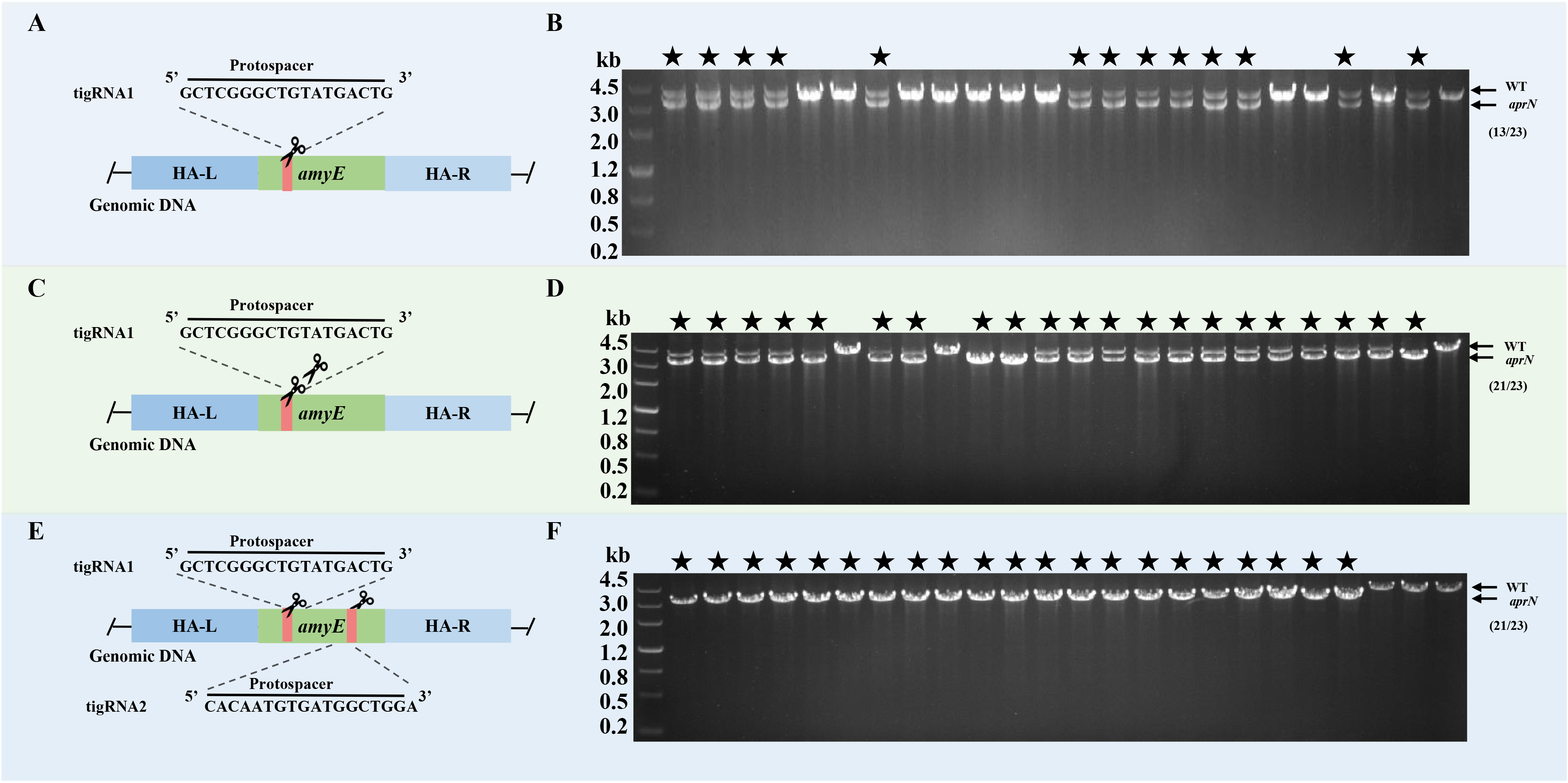
The pBsuTasR system enables gene integration in *Bacillus subtilis* SCK6. (A) Schematic diagram of integration of the *aprN* gene at the amyE site of the SCK6 genome using a single tigRNA. (B) DNA gel image confirming the deletion of a 1146 bp genomic fragment at the same locus mediated by a single tigRNA. (C) Schematic diagram of integration of the *aprN* gene at the amyE site of the SCK6 genome using dual tigRNAs targeting the same locus. (D) DNA gel image confirming the deletion of a 1146 bp genomic fragment at the same locus mediated by dual tigRNAs. (E) Schematic diagram of integration of the *aprN* gene at the amyE site of the SCK6 genome using dual tigRNAs targeting different sites. (F) DNA gel image confirming the deletion of a 1146 bp genomic fragment mediated by dual tigRNAs targeting different sites.

In this study, pBsuTasR-*amyE*-*aprN* was further optimized to address the aforementioned mixed population issue using a dual-tigRNA strategy. This strategy was implemented in two configurations: the first employs two identical tigRNAs to cleave the same position within the target gene (Fig. 4C), termed “same-site dual cleavage”; the second utilizes two distinct tigRNAs to cleave different positions within the same gene (Fig. 4E), termed “different-site dual cleavage”. The results showed that the efficiency of gene integration was improved to 91.3% by using the strategy of “same-site dual cleavage”, but there was still bacteria mixing (Fig. 4D). When the strategy of “different-site dual cleavage” is adopted, the integration efficiency is not only improved to 91.3%, but also the PCR shows that the bands are relatively single (Fig. 4F), which solves the problem of bacteria mixing. These results suggest that employing two distinct tigRNAs to cleave different positions within the same gene represents the optimal strategy for improving editing efficiency to resolve the mixed population issue.

In summary, the integration of *aprN* gene in *amyE* locus of *B. subtilis* SCK6 was successfully realized by TasR in this study. Compared with the previously established enIscB, the single gene integration efficiency of TasR is higher in the same host (enIscB is 6.6% ∼ 26.6%). Compared with CRISPR-Cas9^[31]^ and Cpf1^[14]^, TasR can complete precise integration with a single plasmid, with simpler operation and more efficient process. It provides a new and efficient tool for the stable genome integration and metabolic engineering transformation of *B. subtilis* SCK6.

### 3.5 Sequential gene editing in *B. subtilis* using the “Scissors-Rock-Paper” strategy

The aforementioned experiments confirmed that the pBsuTasR system enables efficient single-locus editing. To meet the demand for sequential iterative editing of multiple genes at multiple loci, and to overcome the technical bottlenecks of cumbersome plasmid elimination and limited selection markers in multi-round genomic editing of *B. subtilis* SCK6, this study designed and constructed the pBsu-SRP iterative genomic editing system adapted for TasR based on the “Scissors-Rock-Paper” strategy. The construction of key elements and functional validation of iterative gene editing are described herein.

The pBsu-SRP system comprises three mutually exclusive editing plasmids: pBsu-Scissors, pBsu-Paper, and pBsu-Rock (Fig. 5A). These three plasmids share an identical replicon and TasR core elements but carry distinct selection markers. Each plasmid expresses a tigRNA that specifically targets the resistance region of another plasmid, establishing a mutual elimination relationship: pBsu-Rock eliminates pBsu-Scissors, pBsu-Paper eliminates pBsu-Rock, and pBsu-Scissors eliminates pBsu-Paper. This design automatically triggers cleavage and removal of the previous round of plasmids upon transformation of a new round, eliminating the need for additional counter-selection or auxotrophic hosts, and providing the core foundation for continuous multi-round genomic editing in SCK6. To validate the feasibility of the designed Scissors-Rock-Paper strategy in *B. subtilis*, the *spo0A*, *amyE*, and *sacB* genes, which had been previously deleted using the pBsu-TasR system in this study, were selected as target loci for functional verification.

**Figure 5.**
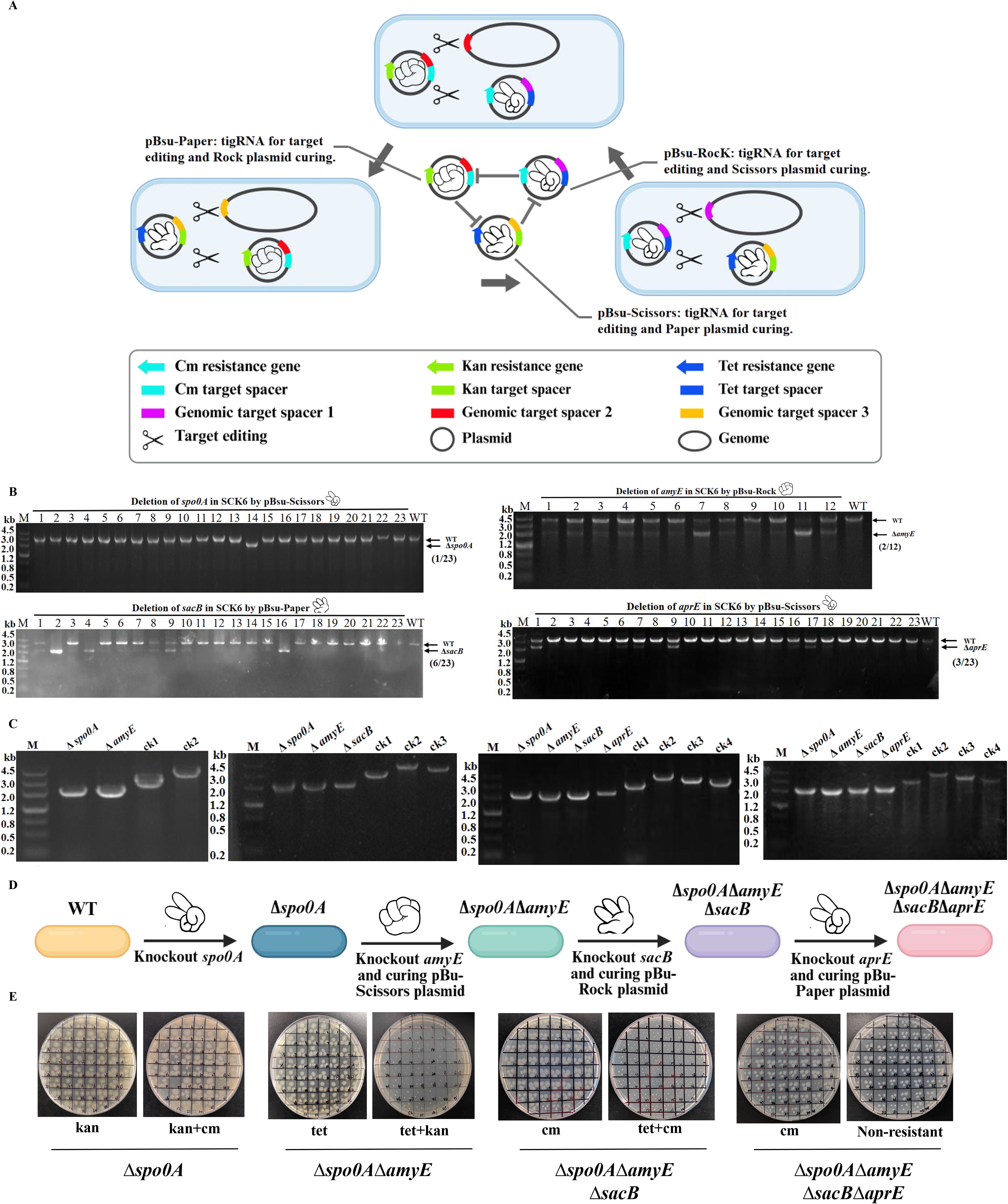
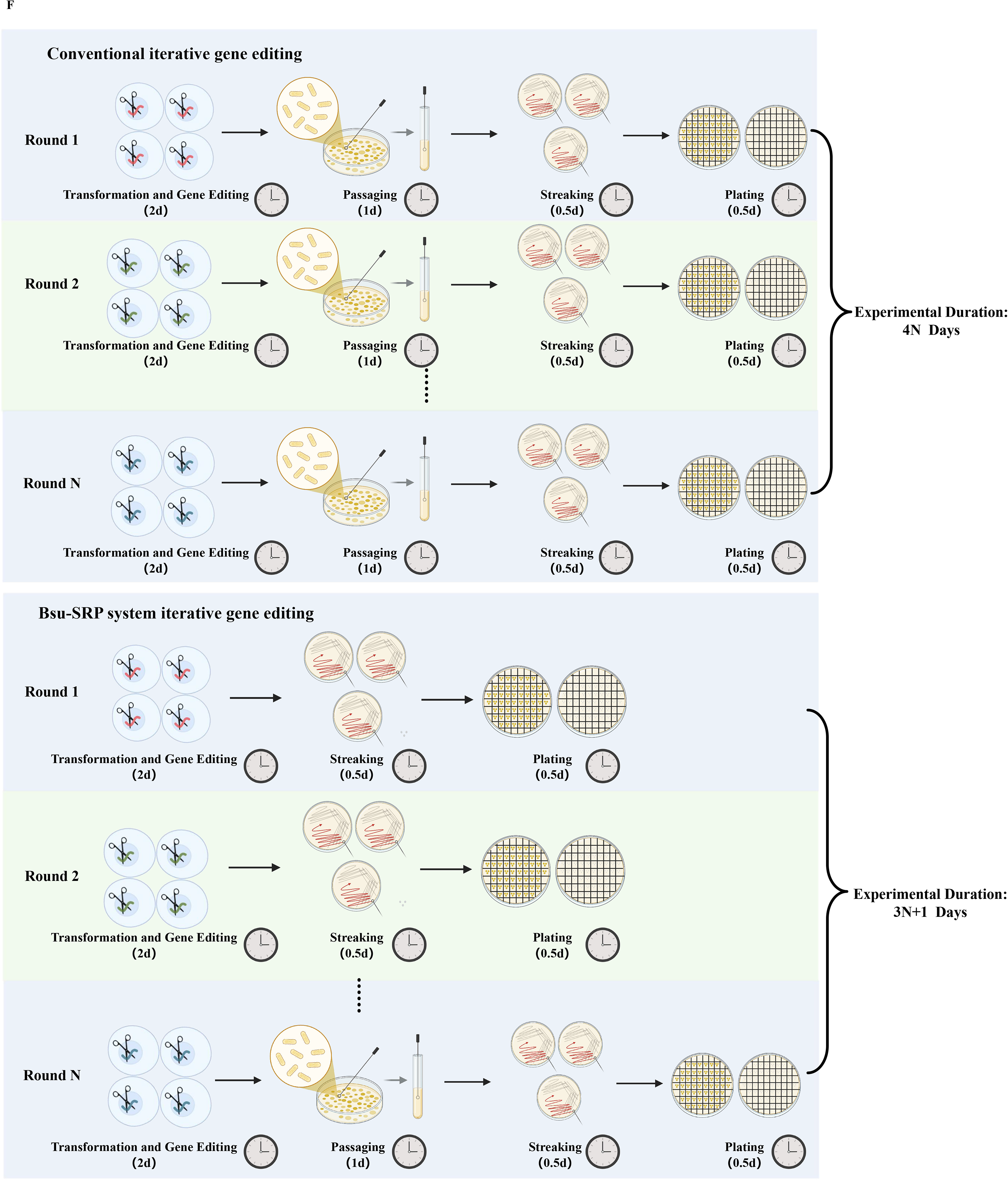
Development and design of key components of the SRP iterative genome editing system in *Bacillus subtilis* SCK6. (A) Schematic diagram of the pBsu-SRP iterative genome editing system principle. (B) DNA gel image sequentially verifying the completion of gene editing by pBsu-Scissors, pBsu-Rock, and pBsu-Paper. (C) DNA gel image verifying the sequential accumulation of *spo0A*, *amyE*, *sacB*, and *aprE* gene deletions after each round of genome editing, with the plasmid from the previous round having been automatically cured. ck1: spo0A WT; ck2: amyE WT; ck3: sacB WT; ck4: aprE WT. (D) Principle of plasmid curing in the SRP iterative genome editing system. (E) Spot plating verification of plasmid curing. (F) Cycle time comparison between the conventional plasmid discard step and the plasmid discard step of the pBsu-SRP system.

First, the *spo0A* locus was targeted for the first round of “Scissors-Rock-Paper” gene editing. The constructed pBsu-Scissors-*spo0A* plasmid was transformed into competent cells of *B. subtilis* SCK6, followed by overnight incubation. A total of 23 transformants were randomly selected for colony PCR verification, with wild-type SCK6 as a control. The results showed that transformant 14 amplified a PCR band of about 2 kb, which was consistent with the expected fragment size after *spo0A* gene deletion, and the deletion efficiency was 4.34% (Fig. 5B). The PCR product was further purified and sequenced, and the results confirmed that *spo0A* gene had been successfully deleted. The above results indicate that the Scissors plasmid pBsu-Scissors enables efficient gene deletion in *B. subtilis*. The *B. subtilis* SCK6 Δ*spo0A* strain was successfully obtained, laying the foundation for subsequent gene editing operations in this study.

Following the successful deletion of *spo0A* in the first round, the *amyE* gene was targeted for the second round of gene editing. The constructed pBsu-Rock-*amyE* plasmid was transformed into competent cells of *B. subtilis* SCK6 Δ*spo0A*, followed by overnight incubation. A total of 12 transformants were randomly selected for colony PCR verification, with wild-type SCK6 as a control. The results showed that 2 out of 12 transformants amplified a PCR band of about 2 kb, which was consistent with the expected fragment size after *amyE* gene deletion, and the deletion efficiency was 16.67% (Fig. 5B). Furthermore, the PCR product was purified and sequenced, and the results confirmed that *amyE* gene had been successfully deleted. Non-specific bands were detected in the colony PCR results, indicating the presence of wild-type strains among the selected colonies. Therefore, transformant No. 11 was subjected to streak purification, and single colonies carrying only the *amyE* deletion were successfully obtained. Genotype verification confirmed the double deletion of *spo0A* and *amyE* (Fig. 5C), yielding the *B. subtilis* SCK6 Δ*spo0A*Δ*amyE* strain.

For the third round of gene editing, the *sacB* gene was targeted as the editing locus. The constructed pBsu-Paper-*sacB* plasmid was transformed into competent cells of *B. subtilis* SCK6Δ*spo0A*Δ*amyE*, followed by overnight incubation. A total of 23 transformants were randomly selected for colony PCR verification, with wild-type SCK6 as a control. The results showed that 6 transformants amplified a PCR band of about 2 kb, which was consistent with the expected fragment size after *sacB* gene deletion, and the deletion efficiency was 26.08% (Fig. 5B). The PCR product was further purified and sequenced, confirming successful deletion of the *sacB* gene. Genotype verification confirmed the triple deletion of *spo0A*, *amyE*, and *sacB* (Fig. 5C), yielding the *B. subtilis* SCK6Δ*spo0A*Δ*amyE*Δ*sacB* strain.

To verify whether pBsu-Scissors could efficiently eliminate the pBsu-Paper plasmid, a fourth round of gene editing was conducted. The *aprE* gene was targeted as the editing locus. The constructed pBsu-Scissors-*aprE* plasmid wastransformed into competent cells of *B. subtilis* SCK6Δ*spo0A*Δ*amyE*Δ*sacB*, followed by overnight incubation. A total of 23 transformants were randomly selected for colony PCR verification, with wild-type SCK6 as a control. The results showed that the PCR band of about 2 kb was amplified by 3 transformants, which was consistent with the expected fragment size after *aprE* gene deletion, and the deletion efficiency was 13.04% (Fig. 5B). The PCR product was further purified and sequenced, and the results confirmed that *aprE* gene had been successfully deleted. Non-specific bands were observed in the colony PCR results, indicating the presence of wild-type strains among the selected colonies. Therefore, transformant No. 1 was subjected to streak purification, and single colonies carrying the *aprE* deletion were successfully obtained. Genotype verification confirmed the quadruple deletion of *spo0A*, *amyE*, *sacB*, and *aprE* (Fig. 5C), yielding the *B. subtilis* SCK6Δ*spo0A*Δ*amyE*Δ*sacB*Δ*aprE* strain.

The pBsu-SRP system enables automatic elimination of the previous-round plasmid via resistance markers carried by the new-round plasmid targeting tigRNA (Fig. 5D). To verify plasmid elimination efficiency, strains successfully edited in each round were subjected to streak dilution, and single colonies were randomly picked for spot plating to confirm whether the editing plasmid from the preceding round had been eliminated. The results showed that the pBsu-Scissors-*spo0A* plasmid was successfully eliminated by the second gene editing strain, and the elimination efficiency was 7.69% (4/52). PBsu-Rock-*amyE* plasmid was successfully eliminated by the third gene editing strain, and the elimination efficiency was 100% (52/52). The fourth round of gene editing strains successfully eliminated the pBsu-Paper-*sacB* plasmid, and the elimination efficiency was 100%(52/52) (Fig. 5E).

Since each editing plasmid carries the temperature-sensitive replicon rep pE194ts for replication in *B. subtilis*, the plasmid can be eliminated by serial passage at 50℃. Using the Scissors-Paper-Rock system, four rounds of continuous gene editing have been completed, yielding the *B. subtilis* SCK6Δ*spo0A*Δ*amyE*Δ*sacB*Δ*aprE* strain with all previous-round editing plasmids eliminated. The next step is to eliminate the pBsu-Scissors-*aprE* plasmid. The correctly edited strain was inoculated into antibiotic-free LB medium and serially passaged in antibiotic-free LB at 50℃, 200 rpm for 8 h per passage for three consecutive passages. The culture was then diluted and plated to obtain single colonies, which were subsequently subjected to chloramphenicol spot testing. A total of 52 strains were selected, and the results showed that No.35 strain was sensitive to chloramphenicol, and no colony grew on chloramphenicol plate, indicating that the editing plasmid pBsu-Scissors-*aprE* had been eliminated (Fig. 5E). Finally, PCR verified the genotype in colony 35, and confirmed that the strain was indeed a four-gene deletion after continuous gene editing (Fig. 5C).

In conventional gene editing workflows, starting from transformation of the editing plasmid into *B. subtilis*, gene editing requires 2 days, followed by plasmid passage (1 day), streak purification (0.5 day), and antibiotic spot testing for verification (0.5 day), totaling 4 days per round. For sequential editing of N genes, the total duration is 4N days. The iterative editing approach based on the “Scissors-Paper-Rock” strategy eliminates the plasmid passage step during editing (saving 1 day), reducing each round to 3 days. After the final round, an additional plasmid passage at 50℃ (1 day) and streak-spot verification (1 day) are required; the streak-spot step is performed both during sequential editing and after the final round. Therefore, the total time for sequential editing of N genes is 3N + 1 days (where “+ 1 day” accounts for the post-final-round plasmid passage). Compared with the conventional method, the pBsu-SRP system reduces the iterative gene editing cycle from 4N days to 3N + 1 days. Consequently, as the number of edited genes increases during strain engineering, the time advantage of this method over the conventional approach becomes increasingly pronounced. In summary, this study successfully constructed and systematically validated the pBsu-SRP iterative genome editing system for *B. subtilis* SCK6, encompassing core component construction, sequential gene editing verification, and automatic plasmid elimination validation. By exploiting the TasR-independent, TAM-free editing advantage and the mutual plasmid elimination mechanism inherent to the pBsu-SRP system, this approach enables efficient sequential gene editing without the need for plasmid passage or counter-selection, establishing a novel technical platform for large-scale metabolic engineering of the SCK6 strain.

### 3.6 Multi-copy gene integration in *B.subtilis*via the “Scissors-Rock-Paper” strategy

Upon completion of the core component construction and preliminary validation of the pBsu-SRP system, *mCherry* was employed as a fluorescent reporter to intuitively and quantitatively characterize the sequential editing capability of this system. Using a *B. subtilis* integration strain with *mCherry* successfully integrated at the *ctc* locus, *epr* and *nprB* were selected as target integration sites. According to the Scissors-Rock-Paper strategy, sequential iterative integration was performed by first integrating a second copy of *mCherry* at the *epr* locus, followed by a third copy at the *nprB* locus (Fig. 6A).

**Figure 6.**
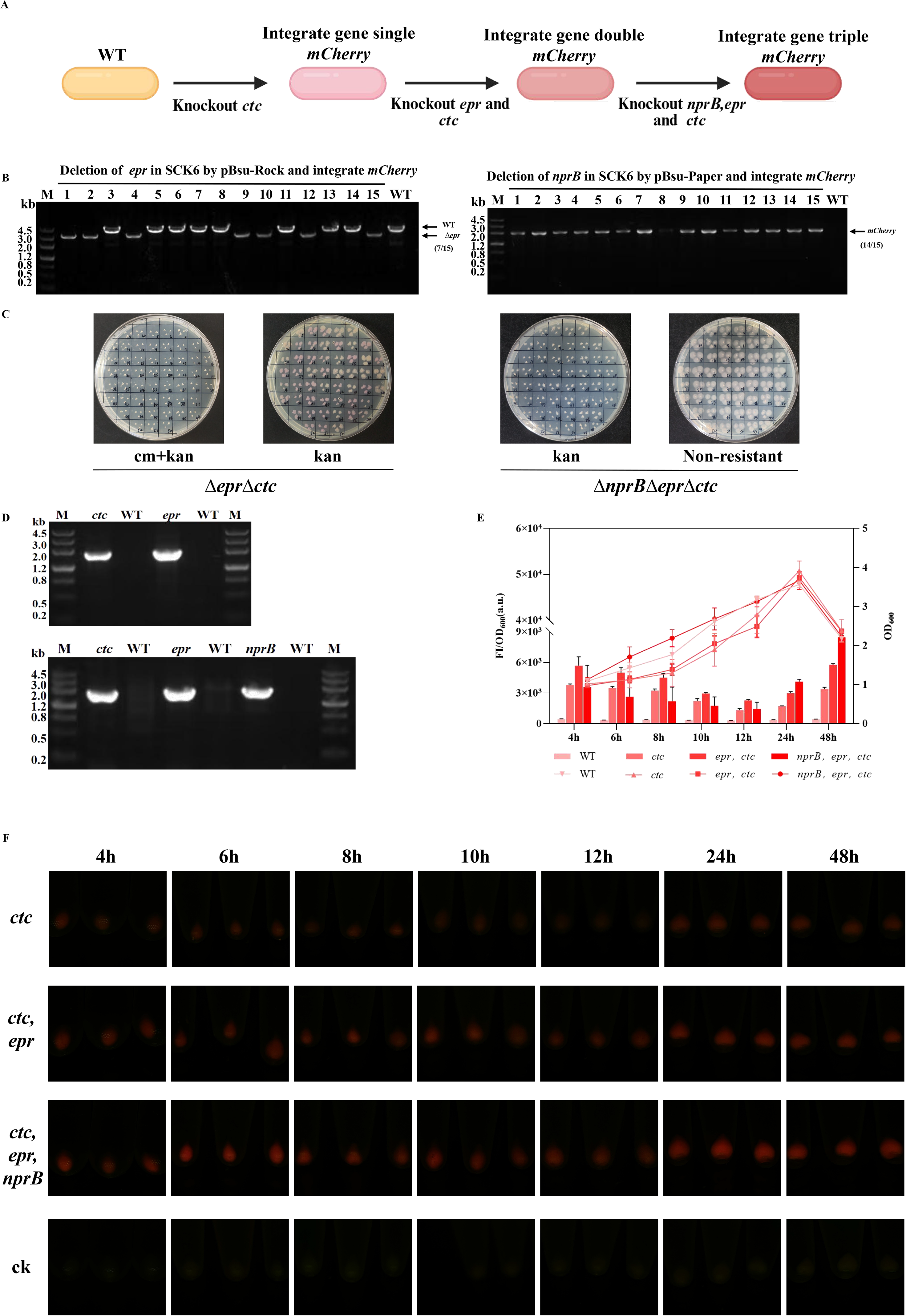
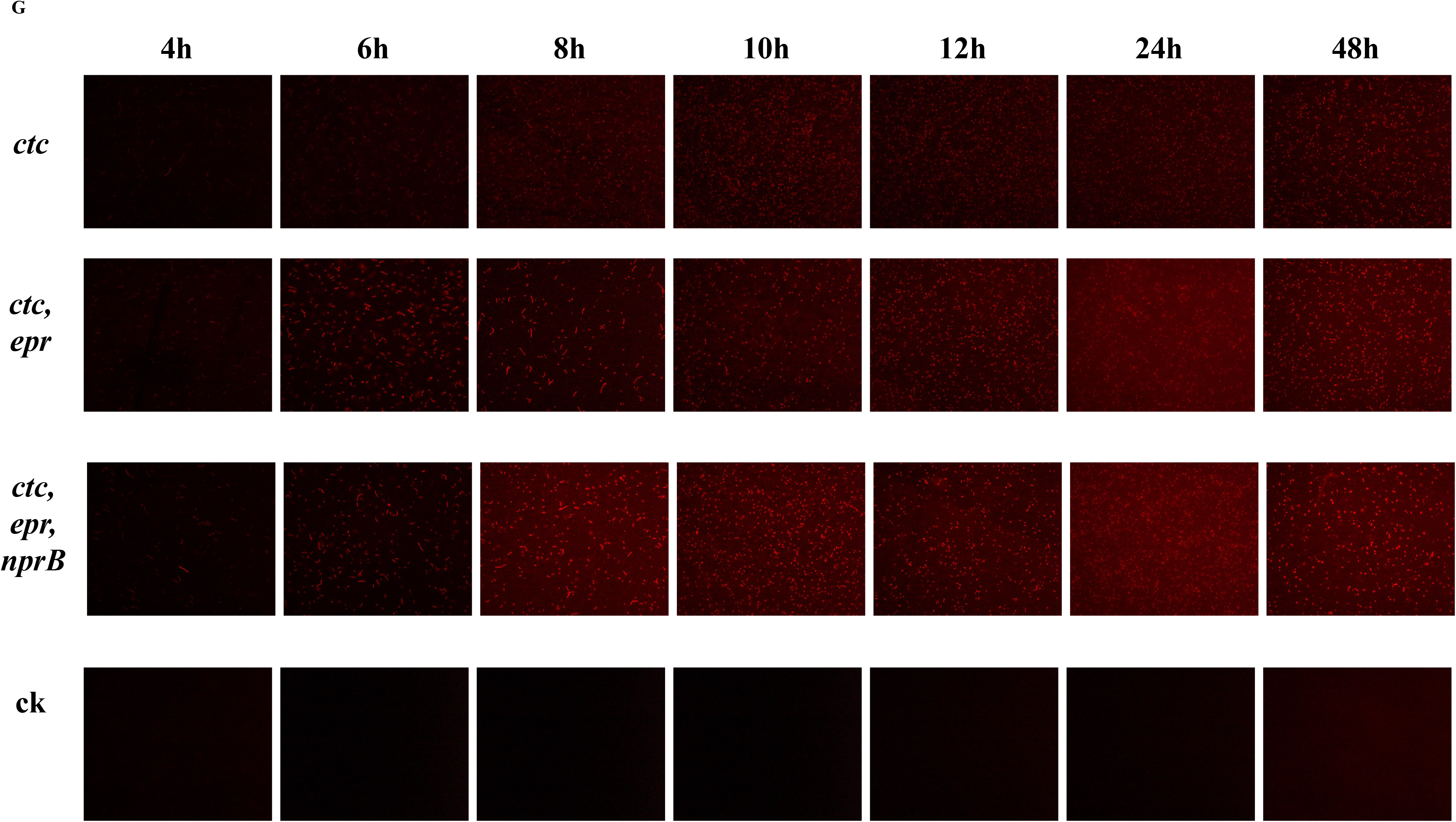
Integration of the *mCherry* gene at different loci of the *Bacillus subtilis* SCK6 genome. (A) Schematic diagram of *mCherry* gene integration at the *ctc*, *epr*, and *nprB* loci based on the principle of the pBsu-SRP system. (B) DNA gel image sequentially verifying the completion of gene editing by pBsu-Scissors-*epr* and pBsu-Rock-*nprB*. (C) Confirmation of plasmid curing in the integrated strains. (D) Genotypes of the integrated strains after plasmid curing. (E) Fluorescence intensity and OD₆₀₀ values of the integrated strains. ck: wild-type SCK6 strain. (F) Macroscopic fluorescence images of bacterial pellets of the integrated strains, observed under a UV transilluminator in the dark. (G) Dynamically captured fluorescence micrographs of the integrated strains obtained from 10 μL of bacterial suspension.

Based on the previously constructed pBsu-Scissors-*spo0A* and pBsu-Rock-*amyE* plasmids, pBsu-Scissors-*epr* and pBsu-Rock-*nprB* were recombinantly constructed and sequentially transformed into *B. subtilis* SCK6 in the order of Scissors-Rock-Paper (SRP). The results showed that the pBsu-SRP system integrated *mCherry* at *epr* and *nprB* sites, and the efficiency was 46.7% and 93.3% respectively (Fig. 6B). The gene integration was verified by sequencing.

This study further investigated the effect of increasing gene copy number on *mCherry* expression levels in *B. subtilis* SCK6. Antibiotic spot testing for plasmid curing was performed on the *mCherry* integration strains obtained at each round. The results showed that, while *mCherry* copies were iteratively integrated round by round, the editing plasmid from the previous round was simultaneously and automatically eliminated (Fig. 6C), and genotypes were confirmed by PCR (Fig. 6D). By dynamically monitoring OD₆₀₀ and fluorescence values, composite curves of fluorescence intensity and bacterial growth were generated. At the same cultivation duration, the fluorescence intensity of the integration strains increased progressively with the number of *mCherry* copies. All integration strains exhibited continuously rising fluorescence levels over time, peaking at 48 h, with the second-highest levels observed at 24 h. In addition, macroscopic fluorescence imaging and fluorescence microscopy were performed on all integration strains at the corresponding time points. The results showed that the wild-type strain exhibited no red fluorescence throughout the entire culture period, whereas the integration strains harboring *mCherry* copies displayed gradually increasing red fluorescence over time. Specifically, the single-copy integration strain began to show weak red fluorescence at 4 h; the double-copy integration strain exhibited higher fluorescence intensity than the single-copy strain at 4 h; and the triple-copy integration strain displayed clearly detectable red fluorescence as early as 4 h, with the highest fluorescence intensity among the three groups across all time points. These results indicate that *mCherry* expression at the *ctc*, *epr*, and *nprB* loci in the integration strains increased within 48 h post-integration, consistent with the fluorometric measurements and OD₆₀₀ values.

In summary, the pBsu-SRP system achieves multi-round marker-free sequential integration through autonomous plasmid elimination. The *mCherry* expression level increases with the number of genomically integrated copies, confirming that the RPS system can efficiently and stably accomplish stepwise, site-specific insertion of multi-copy exogenous genes in *B. subtilis*, demonstrating outstanding application potential for multi-copy gene modular engineering in *B. subtilis* chassis strains.

### 3.7 Escape of *B.subtilis*from TasR cleavage and preliminary investigation of escape patterns

To achieve more thorough cleavage of the *B. subtilis* genome, a dual tigRNA plasmid targeting both ends of the *amyE* gene was designed. The linearized pBsuTasR-*amyE* plasmid backbone and the tigRNA2 fragment were assembled via a one-step cloning method (Fig. 7A). The tigRNA2 was successfully assembled onto the pBsuTasR-*amyE* plasmid template, followed by plasmid extraction from positive *E. coli* colonies. The sequencing results showed that the plasmid sequence was consistent with the expectation, indicating that the plasmid was successfully constructed.

**Figure 7.**
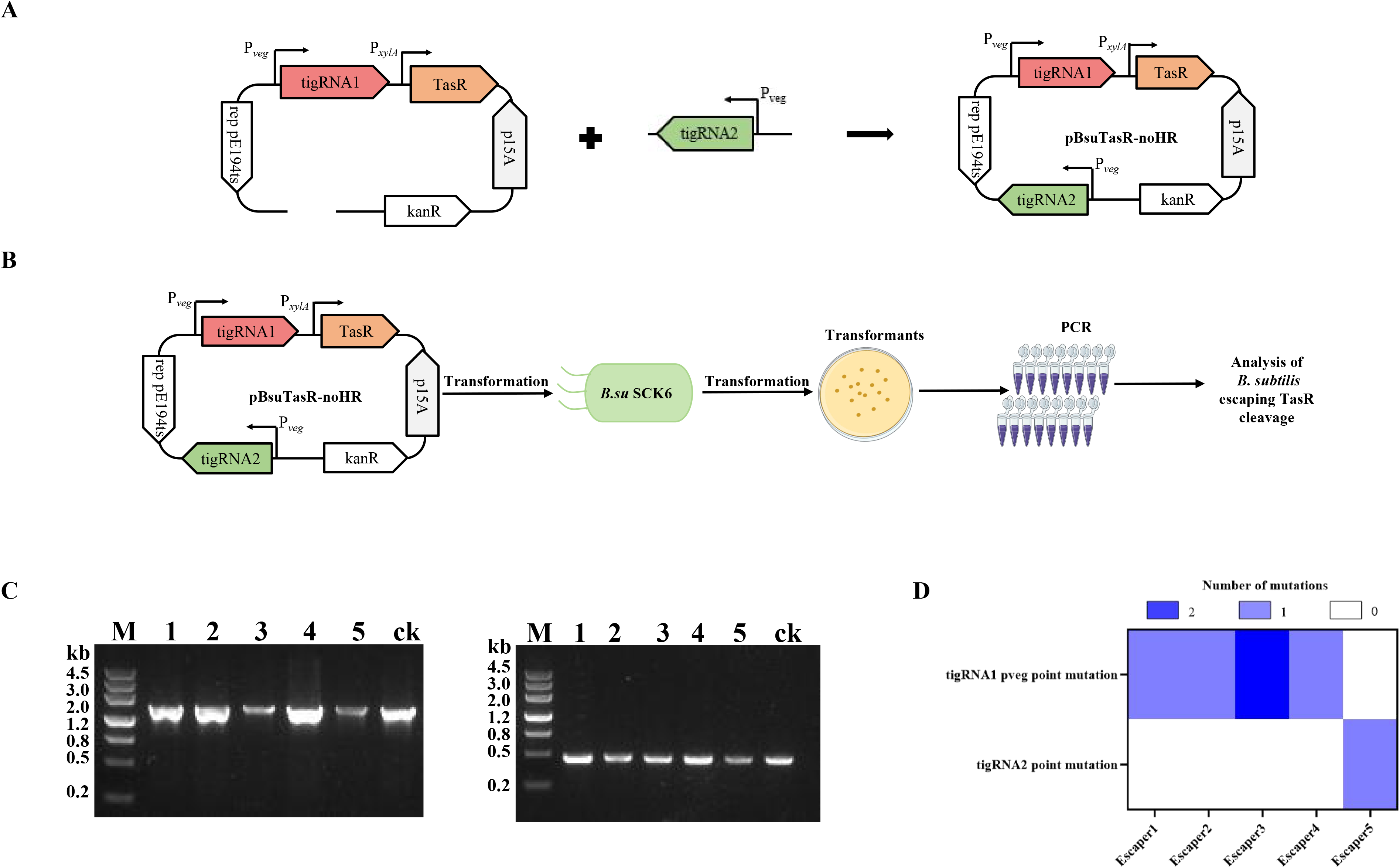
Elucidation of the mechanisms by which *Bacillus subtilis* SCK6 escapes TasR cleavage. (A) Construction workflow of the pBsuTasR-amyE-tigRNA2 plasmid. (B) Workflow of TasR cleavage in SCK6. (C) DNA gel image showing the PCR results of escapers from TasR cleavage in SCK6. (D) Analysis of the mutations in escapers surviving TasR cleavage at the *amyE* site.

The constructed pBsuTasR-*amyE*-tigRNA2 plasmid was transformed into *B. subtilis* SCK6 and plated on LB agar plates supplemented with kanamycin and 3% xylose. After overnight incubation until clearly visible single colonies appeared, these colonies were identified as escape mutants (Fig. 7B). A total of 5 escape mutants were randomly picked, and using the pBsuTasR-*amyE*-tigRNA2 plasmid as a control, the tigRNA1+TasR fragment and the tigRNA2 fragment were amplified by PCR with the corresponding primer pairs. The results showed that the band size was consistent with that of the control, both of which were in line with expectations, and there was no obvious change (Fig. 7C). Therefore, it is necessary to purify the PCR products and sequence them, and further analyze the specific reasons why *B. subtilis* SCK6 escaped DNA nuclease TasR cleavage.

To further investigate the specific mechanisms underlying the escape of *B. subtilis* SCK6 from TasR nuclease cleavage, the amplified fragments from 6 randomly selected escape mutants were purified and subjected to sequencing. The sequencing results were compared with the plasmid map, and the results showed that all 5 escapees had point mutations (Fig. 7D), among which escapees No.1, No.2 and No.4 had 1 bp point mutations at the promoter P_veg_ of tigRNA1, No.3 had 2 bp point mutations at the promoter P_veg_ of tigRNA1, and No.5 had 1 bp point mutations at tigRNA2. From the above results, it can be seen that the point mutation of editing elements is the main reason for *B. subtilis* to escape the cleavage of DNA nuclease TasR, which mainly occurs in the promoter P_veg_ and tigRNA2 of tigRNA1.

In summary, point mutations are the predominant mechanism by which *B. subtilis* SCK6 escapes TasR cleavage, with mutation hotspots primarily concentrated in the tigRNA promoter region and the tigRNA sequence itself. This finding provides an important reference for subsequent optimization of the TasR system, reduction of escape rates, and improvement of editing stability.

### 3.8 Expanding the TasR-based miniature gene editing tool to other bacterial species

Building on our previous successful construction of the pBsuTasR system in *B. subtilis* SCK6, this study aimed to expand the host range of TasR and overcome the inherent TAM sequence dependency and application limitations of the homologous IscB system. To this end, we attempted to construct a TasR-based genome editing system, pEcTasR, in *E. coli* MG1655, and investigated target editing efficiency and the mechanisms underlying escape formation.

First, the *maeB*, *umuDC*, and *lacZ* genes of *E. coli* MG1655 were selected as target genes to verify gene deletion using the pEcTasR system. A total of 15 transformants were randomly picked for colony PCR verification. The results showed that the gene deletion efficiencies of the pEcTasR system were:26.7%(*maeB*), 100%(*umuDC*) and 6.7%(*lacZ*) (Fig. 8A). These results demonstrated that the pEcTasR system was successfully constructed in *E. coli* MG1655. The system exhibited significant variability in editing efficiency across different target genes, with the editing efficiency for the *umuDC* gene reaching 100%, indicating efficient targeted editing capability. DNA sequencing further confirmed that the targeted gene deletion regions in the Δ*maeB*, Δ*umuDC*, and Δ*lacZ* strains were consistent with the expected sequences, with deletion fragment lengths of 2280 bp, 1688 bp, and 3075 bp, respectively (Fig. 8B).

**Figure 8.**
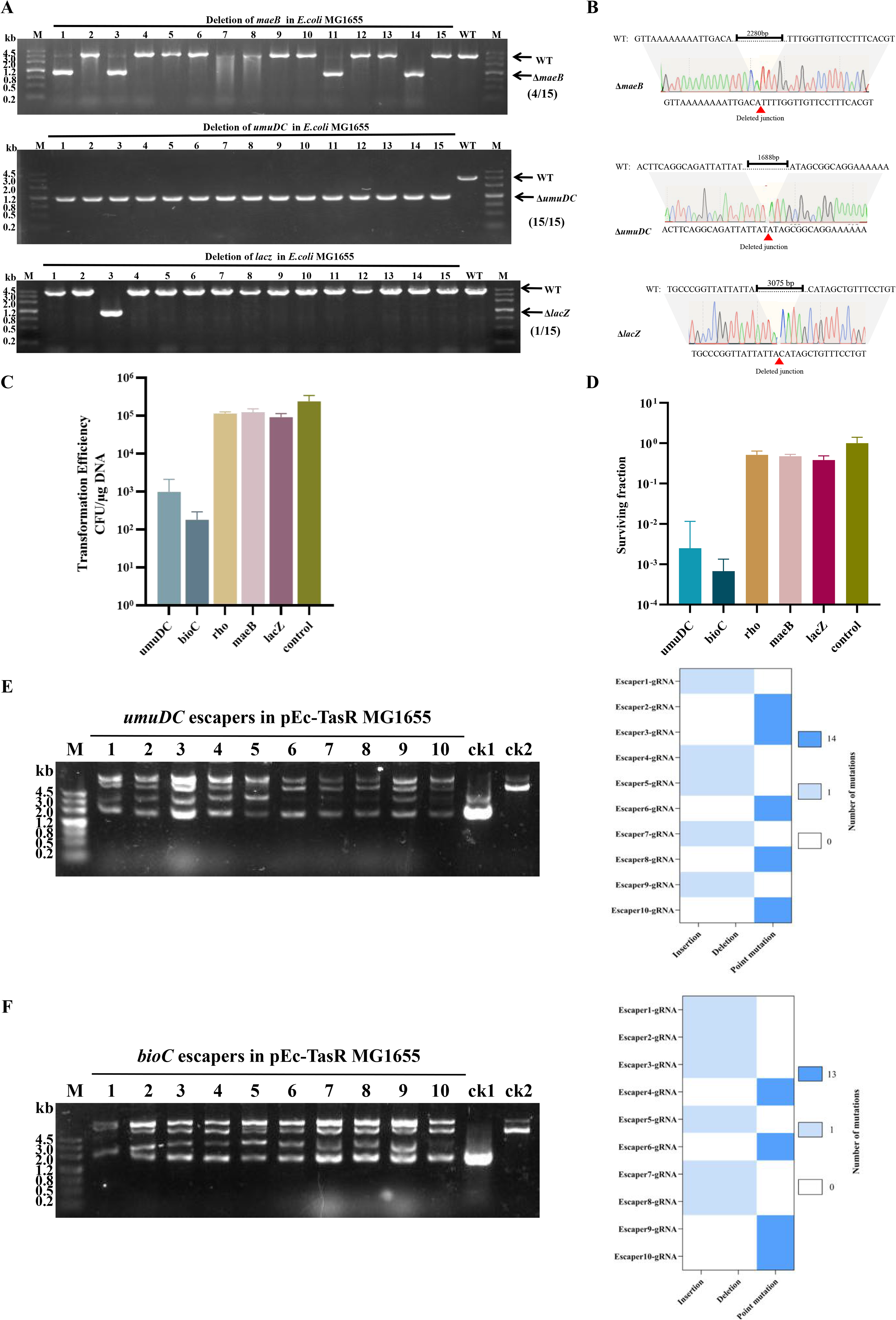
Construction of the pEcTasR system in *Escherichia coli* MG1655 based on the building principle of the pBsuTasR system and its related applications. (A) DNA gel image for PCR verification of *maeB*, *umuDC*, and *lacZ* gene deletions by the pEcTasR system. (B) DNA sequencing results confirming *maeB*, *umuDC*, and *lacZ* gene deletions by the pEcTasR system. Deletion sizes: Δ*maeB*, 2280 bp; Δ*umuDC*, 1688 bp; and Δ*lacZ*, 3075 bp. (C) Transformation efficiency of *E. coli* MG1655 under targeting at different loci. (D) Cell survival fraction of E. coli MG1655 under targeting at different loci. (E) DNA gel image showing the sizes of pEc-TasR and tigRNA series plasmids extracted from escapers obtained at the *umuDC* locus, along with the mutation types and frequencies in the TasR and tigRNA coding sequences of escapers obtained at the *umuDC* locus. ck1, pEc-TasR; ck2, pEc-tigRNA-*umuDC*. (F) DNA gel image showing the sizes of pEc-TasR and tigRNA series plasmids extracted from escapers obtained at the *bioC* locus, along with the mutation types and frequencies in the TasR and tigRNA coding sequences of escapers obtained at the *bioC* locus. ck1, pEc-TasR; ck2, pEc-tigRNA-*bioC*.

Microorganisms can survive RNA-guided DNA nuclease cleavage (termed escapers), and understanding the factors contributing to escape formation can facilitate the development of efficient genome editing tools. To screen for suitable targets for escape mechanism analysis, the plasmid transformation efficiency and strain escape rate were further determined across a total of 5 loci (*umuDC*, *bioC*, *rho*, *maeB*, and *lacZ*), including the aforementioned three genes. The results showed that the transformation efficiency of *umuDC* and *bioC* sites was about 2-3 orders of magnitude lower than that of the control group, indicating that TasR had high cleavage activity for these two sites. However, *rho*, *maeB* and *lacZ* sites decreased slightly and the cutting efficiency was low (Fig. 8C). Further statistics on the escape rate show that the escape rate of *umuDC* and *bioC* sites is much lower than that of *rho*, *maeB* and *lacZ* sites, which is consistent with the cutting efficiency trend (Fig. 8D). Therefore, the non-essential gene *umuDC* and the essential gene *bioC* were selected as the escape target sites for the exploration of *E. coli* MG1655.

Subsequently, tigRNA plasmids targeting the two genes were individually transformed into the *E. coli* MG1655 strain carrying pEcTasR, and escape single colonies were screened and isolated. To investigate the molecular triggers underlying escape formation, 10 escapers were randomly selected from each target locus, and the pEcTasR and tigRNA series plasmids were extracted and examined for size on agarose gels alongside control plasmids (Fig. 7E, F). The results showed that although the pEcTasR plasmid size remained unchanged in the escapers, the tigRNA plasmids at both the *umuDC* and *bioC* loci exhibited significant size alterations, clearly larger than the control CK2. Subsequently, DNA sequencing of the TasR and tigRNA coding regions from 10 escapers targeting the *umuDC* and *bioC* loci revealed multiple mutations.Thermogram analysis showed that 5 escaped strains in *umuDC* group had the same point mutation, and the other 5 had the same deletion/insertion; In the *bioC* group, 4 escaped strains had the same point mutation, and the other 6 had the same deletion/insertion. The characteristics of the two kinds of mutations are highly consistent: point mutations are all located inside tigRNA; The deleted fragments are all 78 nt, which are located inside tigRNA; The inserted fragments are all 161 nt, located at the 5’ end of tigRNA (Fig. 7E and F). These results indicate that mutations in the tigRNA coding sequence are the primary cause of escaper formation in *E. coli* under TasR-mediated genome editing. Further investigation into the mechanisms by which *E. coli* evades TasR editing may facilitate the optimization of TasR-based genome editing tools.

In summary, based on our previous TasR editing work in *B. subtilis*, this study successfully established the pEcTasR editing tool in *E. coli* MG1655 that is independent of TAM sequences, thereby expanding the host range of the TasR protein. The significant effects of target loci on editing efficiency and cell survival provide a reference for subsequent target selection. Furthermore, the elucidation of tigRNA coding sequence mutations as the core escape mechanism offers a theoretical basis for reducing escape rates and improving editing efficiency through tigRNA structural optimization and enhanced stability. Compared with the IscB/TnpB systems reported in the literature, the pEcTasR system demonstrates a significant advantage in target selection flexibility. Additionally, the tigRNA mutation-based escape mechanism identified in this study provides a key theoretical foundation for subsequent site-directed tigRNA modification, enhanced vector stability, reduced escape proportions in engineered strains, and optimized TasR editing efficiency.

## 4 Discussion

*B*, *subtilis* is one of the most widely used industrial microbial chassis in the global biomanufacturing industry, and the efficiency of its genetic modification directly determines the construction timeline and the industrial application potential of high-performance strains^[32]^. Miniature DNA nucleases, by virtue of their small molecular size, provide more flexible vector options for genome editing in *B. subtilis*. However, existing tools still face core challenges, including TAM sequence dependence, strong host specificity, and cumbersome iterative editing workflows^[15,21,33–35]^. In this study, we focused on the TAM-independent miniature nuclease TasR and established a complete toolkit spanning from single-gene editing to successive iterative genome editing in *B*. *subtilis* SCK6. The editing characteristics of TasR were systematically investigated, its escape mechanisms were preliminarily explored, and the system was simultaneously extended to *E*. *coli* MG1655 to verify its universality.

Our research group previously constructed the first IscB-based miniature DNase editing system in *B*. *subtilis* SCK6. However, subsequent experiments revealed that IscB exhibits strict dependence on TAM sequences, which severely restricts its targeting scope. More than half of the genomic genes cannot be efficiently targeted by this tool, making it difficult to fulfill the demands of genome-wide metabolic engineering modification^[15]^. To address this limitation, the present study introduced a novel TAM-independent miniature nuclease, TasR. This protein requires only the formation of a ribonucleoprotein complex with a 36-nt tigRNA to achieve specific DNA cleavage through base pairing, without the need for a TAM sequence, thereby theoretically overcoming the inherent restriction of target selection^[10]^. Based on this property, we constructed the pBsuTasR single-plasmid editing system and successfully achieved the precise deletion of multiple endogenous genes. Compared with the IscB system, the core innovation of TasR lies in completely breaking the constraint imposed by TAM sequences on the editing scope, truly enabling genome-wide targetable editing in *B*. *subtilis*.

After resolving the issue of target coverage for single-gene editing, this study further expanded the editing capability of the TasR system to meet the core demands of genome streamlining and pathway assembly in industrial chassis. In long DNA fragment deletion experiments, we demonstrated that TasR could efficiently mediate the precise deletion of non-essential genomic regions spanning hundreds of kilobases. The optimized dual-tigRNA strategy significantly improved the deletion efficiency, which was comparable to that of our previously developed pBsuenIscB system^[15]^. Compared with the CRISPR-Cas9 two-plasmid system and the Cpf1 system requiring pre-integrated nuclease, pBsuTasR requires only a single transformation to complete the entire procedure, without the need for co-transformation of multiple plasmids or genomic pre-integration steps, thereby greatly shortening the experimental cycle^[14,31]^. Meanwhile, due to its PAM-free nature, optimal cleavage sites can be freely selected at both ends of ultra-large fragments, providing significantly greater flexibility in target design than traditional CRISPR tools. For gene integration, the dual-tigRNA targeting strategy substantially enhanced the site-specific integration efficiency of foreign genes.

However, the construction of complex metabolic pathways often requires multiple rounds of iterative gene editing, and the cumbersome plasmid curing steps in traditional methods have long been a critical bottleneck restricting editing efficiency^[36–39]^. To address this, we innovatively combined the “Scissors-Rock-Paper” strategy with the TasR system to establish the pBsu-SRP iterative genome editing system. By leveraging the targeted cleavage of three mutually exclusive plasmids, this system automatically eliminates the previous-round plasmid upon introduction of the new editing plasmid, without the need for auxotrophic hosts, counter-selectable markers, or additional subculturing throughout the entire process. Using this system, we successfully achieved successive stacked deletions of multiple endogenous genes and verified the feasibility of multi-copy integration via the mCherry fluorescent reporter gene, with the fluorescence expression level increasing as the genomic integration copy number rose. Compared with the conventional workflow in which plasmid curing must be repeatedly performed during multi-round editing, the core advantage of the pBsu-SRP system lies in simplifying the two-step “editing and plasmid curing” procedure into a single transformation, substantially enhancing the efficiency of iterative editing and offering a novel technical route for modular metabolic engineering in *B*. *subtilis*.

While expanding the editing capabilities, this study preliminarily explored the reasons underlying the escape of *B. subtilis* SCK6 from TasR cleavage, which is also a key factor affecting the efficiency of the editing system. Systematic analysis of the escapers revealed that in the TasR editing system, the emergence of escapers was almost exclusively attributable to mutations in the tigRNA coding sequence, and no functional variants of the TasR nuclease itself were detected. In contrast to existing studies, the mechanisms by which *E. coli* and *B. subtilis* escape CRISPR-Cas9 and Cpf1 cleavage are complex and diverse^[40,41]^, encompassing inactivating mutations in the Cas nuclease coding region^[42,43]^, disruption of gRNA expression elements^[43–45]^, as well as target site sequence modifications and target evasion mediated by homologous recombination^[43,46]^. Our group’s previous studies on the escape of *E. coli* from IscB and TnpB cleavage also confirmed that the escape simultaneously involves mutations in the nuclease coding region and gRNA mutations^[21]^. In comparison, the escape mechanism of the TasR system is remarkably singular, with mutations exclusively concentrated in tigRNA and conserved in type. Therefore, subsequent targeted optimization of the tigRNA sequence alone can effectively reduce the escape rate, with a clearly defined optimization direction and substantially lower engineering cost.

To further validate the host universality of the TasR system, we extended it to *E*. *coli* MG1655 and successfully constructed the pEcTasR editing system. This system effectively overcame the strong expression toxicity of IscB in *E. coli* MG1655 and the narrow host range of TnpB, enabling precise deletion of multiple endogenous genes. Further analysis of the *E. coli* escapers revealed that the escape patterns were highly consistent with those observed in *B. subtilis*, with point mutations, fragment deletions, and insertions in the tigRNA coding sequence serving as the primary causative factors. This escape mechanism, highly conserved across Gram-positive and Gram-negative bacteria, not only provides a clear direction for the subsequent design of unified strategies to reduce escape rates but also confirms that the TasR system possesses excellent cross-host adaptability, thereby broadening its application scenarios.

In summary, the TasR editing platform developed in this study combines multiple technical advantages: it retains the convenience of the small molecular size and single-plasmid design characteristic of miniature nucleases, while breaking the target restriction imposed by TAM sequences to achieve genome-wide targetable editing; its long DNA fragment deletion efficiency is comparable to that of mainstream CRISPR tools, yet it offers a simpler operational workflow and greater flexibility in target selection; through the pBsu-SRP system, the challenge of plasmid curing in iterative gene editing has been thoroughly resolved, substantially improving iterative editing efficiency; its escape mechanism is highly singular and conserved, providing a clear direction for subsequent optimization; and the system exhibits excellent cross-host adaptability, functioning effectively in both Gram-positive and Gram-negative bacteria. The technical methodology established in this study has enhanced the genetic modification efficiency of *B*. *subtilis* SCK6, providing a core tool for the rapid construction of high-performance industrial strains capable of producing enzyme preparations and bulk chemicals. This work offers a novel tool for the efficient genetic modification of industrial microbial chassis and will strongly promote the large-scale application of synthetic biology in the green biomanufacturing industry.

## Supporting information

https://fromsmash.com/Supplemental-material

## ASSOCIATED CONTENT

### Supporting Information

### Author Contributions

X.T. and J.G. designed the study, performed key experiments, analyzed data, and drafted the manuscript. H.W., X.W. and X.Z. performed key experiments and assisted with manuscript formatting and literature search. X.P., Y.W. and M.L. provided partial support during the experimental phase. Q.L. conceived and supervised the project, secured funding, provided overall guidance, and critically revised the manuscript.

### Notes

The authors declare no competing financial interest.

## ABBREVIATIONS

FI: fluorescence intensity
GRAS: Generally Recognized as Safe
OD: optical density
SRP: Scissors-Rock-Paper
TAM: transposon-associated motif
tigRNA: TasR-interacting guide RNA

## ACKNOWLEDGEMENTS

This study was supported by National Natural Science Foundation of China (Number: 32402894), Sichuan Science and Technology Program (Number:2024NSFSC0373), and Luzhou Laojiao Co., Ltd (Number: 2022HX07).

