## Supplementary material for "Systematic Development of a Compact Genome-Editing Tool Leveraging the TAM-Independent DNA Nuclease TasR in *Bacillus subtilis*": https://fromsmash.com/Supplemental-material

Table S1: Plasmids and strains used in this study

| Strains or plasmids | Description | Source |
| --- | --- | --- |
| Strains |  |  |
| E. coli DH5α | Commercial transformation host | GIBCO BRL, life Technologies |
| E.coli MG1655 | Host for testing the genome editing efficiency | Lab storage |
| B. su SCK6 | Host for testing the genome editing efficiency | Lab storage |
| Plasmids |  |  |
| pHT-XCR6 |  | [1] |
| pcrF19-NM2 |  | [1] |
| phy300plk |  | [1] |
| pBsu-TasR | Derived from pIscB, inducibly expressing TasR | This study |
| pEc-TasR | Derived from p15A, inducibly expressing TasR | This study |
| pBsuTasR-tigRNA-spo0A | Derived from pBsu-TasR, target spo0A gene |  |
| pBsuTasR-tigRNA-sacB | Derived from pBsu-TasR, target sacB gene |  |
| pBsuTasR-tigRNA-amyE | Derived from pBsu-TasR, target amyE gene |  |
| pBsuTasR-spo0A | Derived from pBsuTasR-tigRNA-spo0A, carry the homology arms of spo0A | This study |
| pBsuTasR-sacB | Derived from pBsuTasR-tigRNA-sacB, carry the homology arms of sacB | This study |
| pBsuTasR-amyE | Derived from pBsuTasR-tigRNA-amyE, carry the homology arms of amyE | This study |
| pBsuTasR-tigRNA-169.9-1 | Derived from pBsu-TasR, target 169.9 genomic fragment |  |
| pBsuTasR-tigRNA-169.9-2 | Derived from pBsu-TasR, target 169.9 genomic fragment |  |
| pBsuTasR-169.9-1 | Derived from pBsuTasR-tigRNA-169-1, carry the homology arms of 169.9 genomic fragment | This study |
| pBsuTasR-169.9-2 | Derived from pBsuTasR-tigRNA-169-2, carry the homology arms of 169.9 genomic fragment | This study |
| pBsuTasR-amyE-aprN | Derived from pBsuTasR-tigRNA-amyE, carry the homology arms of amyE | This study |
| pBsuTasR-amyE-aprN-1 | Derived from pBsuTasR-tigRNA-amyE, carry the homology arms of amyE | This study |
| pBsuTasR-amyE-aprN-2 | Derived from pBsuTasR-tigRNA-amyE, carry the homology arms of amyE | This study |
| pBsu-Scissors-spo0A | Derived from pBsuTasR-tigRNA-spo0A and pHT-XCR6, carry the homology arms of spo0A | This study |
| pBsu-Rock-amyE | Derived from pBsuTasR-tigRNA-amyE, carry the homology arms of amyE | This study |
| pBsu-Paper-sacB | Derived from pBsuTasR-tigRNA-sacB, pHT-XCR6 and phy300plk, carry the homology arms of sacB | This study |
| pBsu-Scissors-aprE | Derived from pBsu-Scissors-spo0A, carry the homology arms of aprE | This study |
| pBsu-Scissors-epr | Derived frompBsu-Scissors-spo0A, carry the homology arms of epr | This study |
| pBsu-Rock-nprB | Derived from pBsu-Rock-amyE, carry the homology arms of nprB | This study |
| pBsuTasR-amyE-tigRNA2 | Derived from pBsuTasR-amyE, target amyE gene | This study |
| pEc-tigRNA-umuDC | Derived from pEc-TasR, target umuDC gene | This study |
| pEc-tigRNA-maeB | Derived from pEc-TasR, target maeB gene | This study |
| pEc-tigRNA-lacZ | Derived from pEc-TasR, target lacZ gene | This study |
| pEc-tigRNA-bioC | Derived from pEc-TasR, target bioC gene | This study |
| pEc-tigRNA-rho | Derived from pEc-TasR, target rho gene | This study |

Table S2: Oligonucleotides used in this study.

| Oligos | Sequence (5’→3’) |
| --- | --- |
| 169-ko-ce-dn | aagattacaatggagtggag |
| 169-pur-up | gacatgaacatcatcatcaatc |
| 16s-q-dn1 | atgcaccacctgtcactctg |
| 16s-q-dn2 | attaccgcggctgctgg |
| 16s-q-up1 | ttcgaagcaacgcgaagaac |
| 16s-q-up2 | actcctacgggaggcagcag |
| Amp-dn | cgttaagggattttggtcattaccaatgcttaatcagtga |
| Amp-gj-dn | aataaacaaataggggttccgcgtggccattcgccattct |
| Amp-gj-up | tcactgattaagcattggtaatgaccaaaatcccttaacg |
| Amp-up | gtgtacagaatggcgaatggccacgcggaacccctatttg |
| amyE-ce-N18-up | gtcatcaatcataccaccag |
| amyE-dndn2 | ttgaacgatgacctcgagcccacgatgcgatttgtgccgatgtacacgtcatctgc |
| amyE-dn-pveg | catttttgtcaaataaaatttaaattatatcaacgttaataaaccgatgtgaagactgg |
| amyE-dnup2 | ctccagtcttcacatcggtgggcaaggctagacggg |
| amyE-HR-dndn | cgatgtacacgtcatctgc |
| amyE-HR-dnup | ctgctcattacactgcgggctctttttcatggtgggcaaggctagacggg |
| amyE-HR-gj-dn | gtcgggttcacacagttgacgactctgagagatcccctcataatttcagc |
| amyE-HR-gj-up | acgggcaatacggtgcagatgacgtgtacatcggctcgaggtcatcgttc |
| amyE-HR-updn | tcttgacactccttatttgat |
| amyE-HR-upup | gtcgtcaactgtgtgaac |
| amyE-in-ce-up | gatttgtattcactctgcc |
| amyE-ko-dn | agttcagctcagtgatacc |
| amyE-ko-dn2 | gctgtgaaggaactgttc |
| amyE-ko-up | gaatcacgagacaggttgc |
| amyE-ko-up2 | ctgatttgatgggaatcacg |
| amyE-pur-dn | tagtgggagatattcgacac |
| amyE-updn2 | cccgtctagccttgcccaccgatgtgaagactggag |
| aprE-dndn-cm | tgacctcgagctagcgacggagagttaggcaaatcctctgttgatatggt |
| aprE-dnup | gaacctgcttctttttactatctttaccctctccttttaa |
| aprE-KO-ce-dn | cacgcaggtcatttgaacg |
| aprE-ko-ce-up | ggatcgagttgacagagaac |
| aprE-ko-dn | taatgccggaataagcaac |
| aprE-ko-dn2 | tgacgatattgcctcctgc |
| aprE-ko-up | caagaaatggatcgaagcg |
| aprE-pur-up | ccaagatatgttgcagtgc |
| aprE-tigRNA-dn | gcgcaaccggctactgggtttcaacgattgcctggctacatttattgtacaacacgagc |
| aprE-tigRNA-up | atgtagccaggcaatcgttgaaacccagtagccggttgcgcaagcaagcttaaacccag |
| aprE-updn | ttaaaaggagagggtaaagatagtaaaaagaagcaggttcct |
| aprE-upup | aaattatgaggggatctctcagagtgctttcgctgattac |
| aprn-ce-up | cagcggaattgactcttc |
| aprN-dn | ttattgtgcagctgcttg |
| aprn-gj-up2 | tatggaaaagggttaatcaacgtacaagcagctgcacaataaagagtcgacctgttacg |
| aprN-up-pveg | ttttatttgacaaaaatgggctcgtgttgtacaataaatgtgtgagaagcaaaaaattg |
| bra-his-dn | acaggtcgactcttcagtggtggtggtggtggtgctcgag |
| bra-his-dn2 | aggtcgactcttcagtggtggtggtggtggtgatattcac |
| bra-his-gj-up | agcaccaccaccaccaccactgaagagtcgacctgttacg |
| bra-his-gj-up2 | atcaccaccaccaccaccactgaagagtcgacctgttacg |
| cm-dn | gatataatgggtttaatagctgaataagaacggtgctcgcaatagttacccttattatc |
| cm-dn-new | aaattatctgaaaggggaatgaactttaataaaattgatttagac |
| cm-gj-dn | ttataaacgatgacctcgagctagcgacggagagttaggagacttactgatcaaacgtg |
| cm-gj-up | caattttattaaagttcattcccctttcagataattttag |
| cm-up | agtctcctaactctccgtcgctagctcgaggtcatcgtttataaaagccagtcattagg |
| cm-up-aprE | accatatcaacagaggatttgcctaactctccgtcgctagctcgaggtcatcgtttata |
| cm-up-ctc | ctgttctcgcaggaaaccctaactctccgtcgctagctcgaggtcatcgtttataaa |
| cm-up-epr | tatcccaatgatgggcctaactctccgtcgctagctcgaggtcatcgtttataaa |
| cm-verf-dn | tatttgaaccaacaaacgac |
| cm-verf-up | agtcattaggcctatctgac |
| ctc-dndn-cm | tataaacgatgacctcgagctagcgacggagagttagggtttcctgcgagaacag |
| ctc-updn-pveg | caaataaaatttaaattatatcaacgttaataaattcagcaccatcctcttg |
| dTasR-dn | agaccaagcgaccgccaatattgtgatatc |
| dTasR-up | atatcacaatattggcggtcgcttggtctcatgaagaacg |
| epr-dndn-cm | tgacctcgagctagcgacggagagttaggcccatcattgggatatgaagc |
| epr-tigRNA-dn | gcgcagagccaagctgggtttcacgataatcctggctacatttattgtacaacacgagc |
| epr-tigRNA-up | atgtagccaggattatcgtgaaacccagcttggctctgcgcaagcaagcttaaacccag |
| kan-dn | gtgcttgaaatcccctcaaaaacc |
| kan-gj-up | atcgggtttttgaggggatttcaagcacaaatcgcatcgtgggactctagaggcactgg |
| kan-up | gcacaaatcgcatcgtgggctcgaggtcatcgttc |
| kc-HR-gj-aprE-dn | gtaatcagcgaaagcactctgagagatcccctcataattt |
| kc-spo0A-ko-dn | gccccatttattcaaaaggc |
| kc-tigRNA2-dn | gactctagaggcactggc |
| kc-tigRNA2-gj-dn | tggccagtgcctctagagtcttgaaatcccctcaaaaacc |
| kc-TnpB-verf-dn | accatgaaaaagagcccg |
| kcTnpB-verf-up | acaatggacacctagggg |
| ko-spo0A-vf-dn | gccccatttattcaaaaggc |
| mcherry q dn3 | gtcctcgaagttcatcacgc |
| mcherry q up3 | caagctgaaggtgaccaagg |
| mecherry-up | ttacttgtacagctcgtccatg |
| no-HR-dn | tttgaacgatgacctcgagctctgagagatcccctcataa |
| no-HR-up | ttatgaggggatctctcagagctcgaggtcatcgttc |
| nprB-dndn-2 | gaacgatgacctcgagcccacgatgcgatttgtgcgtaccatggcatgttcc |
| nprB-ko-dn | gccgatgatgacgatgag |
| nprB-ko-up | ggagaaatacctggaaacc |
| nprB-up-pveg | aatgggctcgtgttgtacaataaatgtatgcgcaacttgaccaag |
| pcrf19-pveg-dn | catttttgtcaaataaaatttaaattatatcaacgttaataacgcatcacacgcaaaaa |
| pcrf19-pveg-up | ttgaggggatttcaactgcagacacctaaattcaaaatctatcggtcag |
| pcrf19-repB-dn | catttttgtcaaataaaatttaaattatatcaacgttaataacgcatcacacgcaaaaa |
| pg-gj-dn | agattttgaatttaggtgtctgcagttgaaatcccctcaa |
| pg-gj-dn-rpeB | tgcagttgaaatcccctcaa |
| pveg-amyE2.2-opup | gcgcagtatgactgtgggtttcaagcccgagctggctacatttattgtacaacacgagc |
| pveg-dn-2 | ttattaacgttgatataatttaaattttatttg |
| pveg-up | aatttaaattttatttgacaaaaatgggctcgtgttgtac |
| repB-up | ttgaggggatttcaactgcactactctttaataaaataatttttccgttccc |
| rep-ce-up1 | gtatccctatcatgtaatgg |
| sacB-dndn | gaacgatgacctcgagcctaatggatctgtgcttttgc |
| sacB-dndn-tet | ctaaacgatgacctcgagcctgcatttccagcactcgctaatggatctgtgcttttgc |
| sacB-dnup | ataaaaaaggagacatgaacgaaacgcaaaagaaaatgcc |
| sacB-HR-gj-dn | gaaggctggatgtacagctctgagagatcccctcataatttcagc |
| sacB-HR-gj-up | cagcaaaagcacagatccattaggctcgaggtcatcgttc |
| sacB-ko-ce-up | gataaagcaggcaagacc |
| sacB-ko-dn | ccaattgttcatccaggc |
| sacB-ko-up | catagctcatgccagatgc |
| sacB-pur-dn | ctgtgttagatgcaatcagc |
| sacB-updn | ggcattttcttttgcgtttcgttcatgtctccttttttatgtac |
| sacB-upup | aattatgaggggatctctcagagctgtacatccagccttc |
| spl-up-amyE | gtcttcaaaaaatcaaataaggagtgtcaagattaaataccaaaaatgtgttgac |
| spo0A-ko-ce-up | gacacatctagggtcgatc |
| spoA-ko-ce-up | gacacatctagggtcgatc |
| Ta-gj-dn-spo0A | gcgcaccgacaaggtgggtttcaccatatccgtggctacatttattgtacaacacgagc |
| TasR-ce-up | tctaaagtgatttgggcgt |
| TasR-gj-dn-amyE | gcgcaatggctggatgggtttcacacattgtgtggctacatttattgtacaacacgagc |
| TasR-gRNA-amyE-up | tgtagccacacaatgtgtgaaacccatccagccattgcgcaagcaagcttaaacccag |
| Ta-ter-up-spo0A | atgtagccacggatatggtgaaacccaccttgtcggtgcgcaagcaagcttaaacccag |
| tet-ce-dn1 | ttgaacgtctcattacctg |
| tet-ce-dn2 | actgccgaaatcggaagtg |
| tet-dn | tgcctctagagtcctgcatttccagcactcgttcaacaaacgggccatat |
| tet-gj-dn | ctaaacgatgacctcgagcctgcatttccagcactcgttcaggtgaaagtgacag |
| tet-gj-up | ggcccgtttgttgaacgagtgctggaaatgcaggactctagaggcactgg |
| tet-up | ctgtcactttcacctgaacgagtgctggaaatgcaggctcgaggtcatcgtttag |
| tet-up-new | acctgaacgagtgctggaaatgcaggctcgaggtcatcgtttagaaatccctttgagaa |
| tet-up-sacB | aaagcacagatccattagcgagtgctggaaatgcaggctcgaggtcatcgtttag |
| tet-up-sacB2 | ccattagcgagtgctggaaatgcaggctcgaggtcatcgtttagaaatccctttgagaa |
| thrC-dndn-kan | ttgaacgatgacctcgagcccacgatgcgatttgtgcgtcgtcatcgaggtgatc |
| thrC-tigRNA-dn | cttgcgcatcgaaggcatgggtttcaagtgtgcgatggctacatttattgtacaacacg |
| thrC-tigRNA-dn3 | cttgcgcagctacgacctgggtttcatgtgttaactggctacatttattgtacaacacg |
| thrC-tigRNA-up | atgtagccatcgcacacttgaaacccatgccttcgatgcgcaagcaagcttaaacccag |
| thrC-tigRNA-up3 | atgtagccagttaacacatgaaacccaggtcgtagctgcgcaagcaagcttaaacccag |
| tig-kan-dn | agccagcacaaatctgaaacccaccacgatgctgcgcaagcaagcttaaacccag |
| tig-kan-up | cagcatcgtggtgggtttcagatttgtgctggctacatttattgtacaacacgag |
| tig-mcherry-vf-dn | tgggtttcacaccatacg |
| tigNA-cmN18-up2 | tattgcgagcaccgttcttattcagctattaaacccattatatcgggtttttgagggga |
| tig-nprB-dn | cttgcgcaacgacggtgtgggtttcagtggatagatggctacatttattgtacaacacg |
| tig-nprB-up | agccatctatccactgaaacccacaccgtcgttgcgcaagcaagcttaaacccagctca |
| tigRNA-169.1-dn | gcgcacgagaaccttgggtttcagcgaagctgtggctacatttattgtacaacacgagc |
| tigRNA-169.1-up | atgtagccacagcttcgctgaaacccaaggttctcgtgcgcaagcaagcttaaacccag |
| tigRNA-169.2-dn | gcgcactgtcagcgtgggtttcaaaggagtcgtggctacatttattgtacaacacgagc |
| tigRNA-169.2-up | atgtagccacgactcctttgaaacccacgctgacagtgcgcaagcaagcttaaacccag |
| tigRNA-169-dn | gcgcaccatcagtttgggtttcagacagctcatggctacatttattgtacaacacgagc |
| tigRNA-169-dn2 | cgcactacctcgatgggtttcacaagtgagctggctacatttattgtacaacacgagc |
| tigRNA-169-up | agccatgagctgtctgaaacccaaactgatggtgcgcaagcaagcttaaacccag |
| tigRNA-169-up2 | agccagctcacttgtgaaacccatcgaggtagtgcgcaagcaagcttaaacccag |
| tigRNA-169-verf-dn | tgggtttcagacagctca |
| tigRNA2-gj-up | atttttgtcaaataaaatttaaattatatcaacgttaataactgcagacacctaaattc |
| tigRNA2-gj-up2 | caaataaaatttaaattatatcaacgttaataagagcaccgttcttattc |
| tigRNA2-verf-dn | gatgaacaaggaatcgtag |
| tigRNA-amyE2.2-opdn | atgtagccagctcgggcttgaaacccacagtcatactgcgcaagcaagcttaaacccag |
| tigRNA-cm-dn | atgtagccacctaactcttgaaacccatagcgacggtgcgcaagcaagcttaaacccag |
| tigRNA-cmN18-dn | tagagtctagcgacggagagttaggttgaaatcccctcaaaaacc |
| tigRNA-cmN18-up | atcgggtttttgaggggatttcaacctaactctccgtcgctagactctagaggcactgg |
| tigRNA-cm-up | cttgcgcaccgtcgctatgggtttcaagagttaggtggctacatttattgtacaacacg |
| tigRNA-ctc-dn | gcgcaagccattggtgggtttcatctccggtctggctacatttattgtacaacacgagc |
| tigRNA-ctc-up | atgtagccagaccggagatgaaacccaccaatggcttgcgcaagcaagcttaaacccag |
| tigRNA-mcherry-dn | gcgcacgtatggtgtgggtttcacaccatacgtggctacatttattgtacaacacgagc |
| tigRNA-mcherry-dn2 | gcgcaagcaagggctgggtttcacaccatacgtggctacatttattgtacaacacgagc |
| tigRNA-mcherry-up | atgtagccacgtatggtgtgaaacccacaccatacgtgcgcaagcaagcttaaacccag |
| tigRNA-mcherry-up2 | atgtagccacgtatggtgtgaaacccagcccttgcttgcgcaagcaagcttaaacccag |
| tigRNA-sacB-dn | gcgcacgacaaccatgggtttcacctgagctgtggctacatttattgtacaacacgagc |
| tigRNA-sacB-dn2 | gcgcaatatgctgctgggtttcacatggcgtgtggctacatttattgtacaacacgagc |
| tigRNA-sacB-up | atgtagccacagctcaggtgaaacccatggttgtcgtgcgcaagcaagcttaaacccag |
| tigRNA-sacB-up2 | atgtagccacacgccatgtgaaacccagcagcatattgcgcaagcaagcttaaacccag |
| tigRNA-spl-dn | gcgcaaaccagcagtgggtttcagtcaacacatggctacatttattgtacaacacgagc |
| tigRNA-spl-up | atgtagccatgtgttgactgaaacccactgctggtttgcgcaagcaagcttaaacccag |
| tigRNA-tet-dn | atgtagccacgagtgctgtgaaacccactgcatttctgcgcaagcaagcttaaacccag |
| tigRNA-tet-up | gcgcagaaatgcagtgggtttcacagcactcgtggctacatttattgtacaacacgagc |
| tigRNA-thrC-dn2 | cttgcgcattacagctttgggtttcaaggcagccgtggctacatttattgtacaacacg |
| tigRNA-thrC-up2 | atgtagccacggctgccttgaaacccaaagctgtaatgcgcaagcaagcttaaacccag |
| tigRNA-verf-up | tccattcactattctcattcc |
| tig-spl-vf-dn | tgggtttcagtcaacaca |

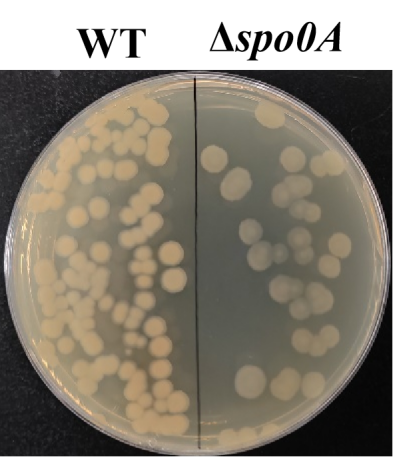

Fig. S1. Phenotypic verification of the knockout *B. subtilis* strain SCK6Δ*spo0A*.

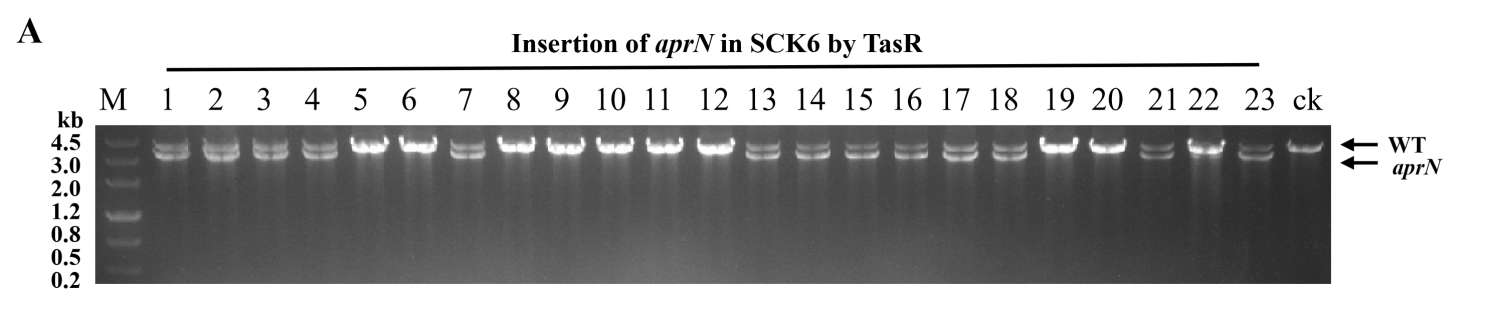

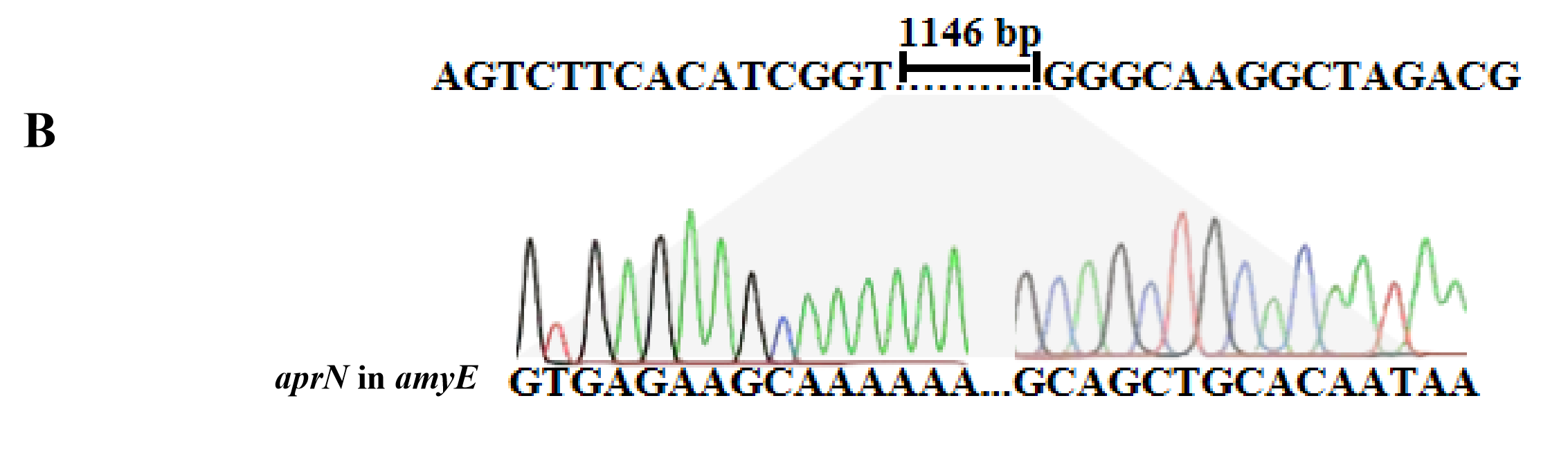

Fig. S2. (A) PCR validation results of aprN gene integration using the pBsuTasR system; (B) DNA sequencing results of aprN gene integration using the pBsuTasR system.

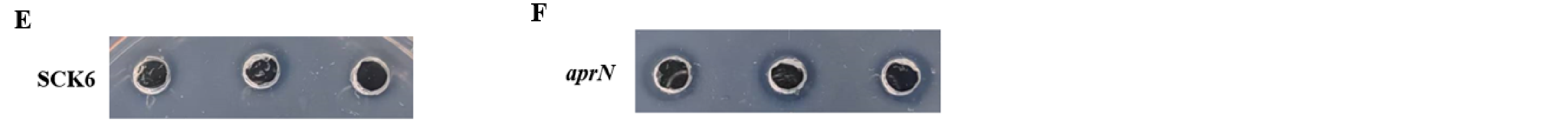

Fig. S3. SCK6:Phenotypic validation of nattokinase activity in the wild-type SCK6 strain; *aprN*:Phenotypic validation of nattokinase activity in the *aprN* gene-integrated strain after plasmid elimination. The black asterisk indicates successful integration.

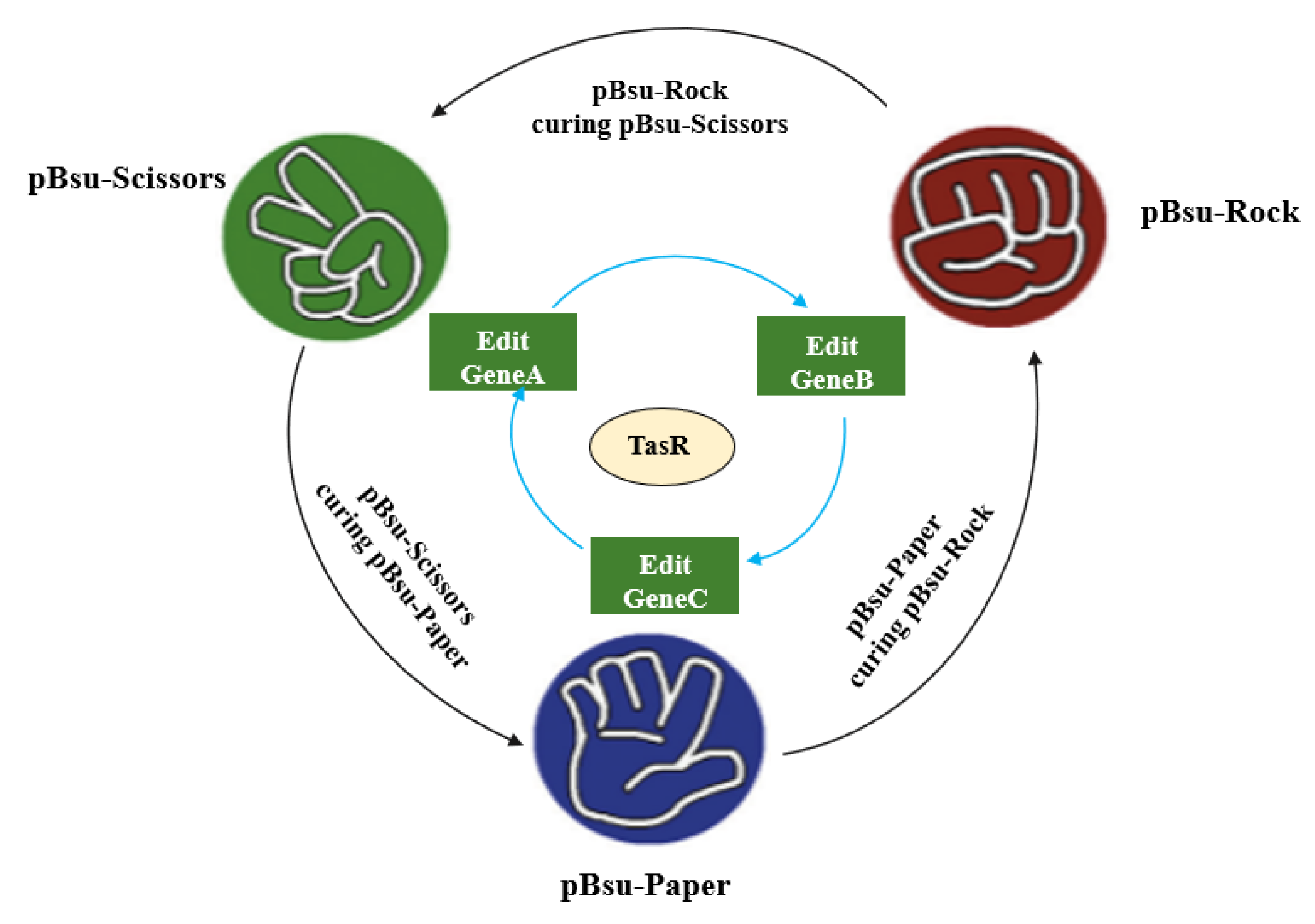

**A**

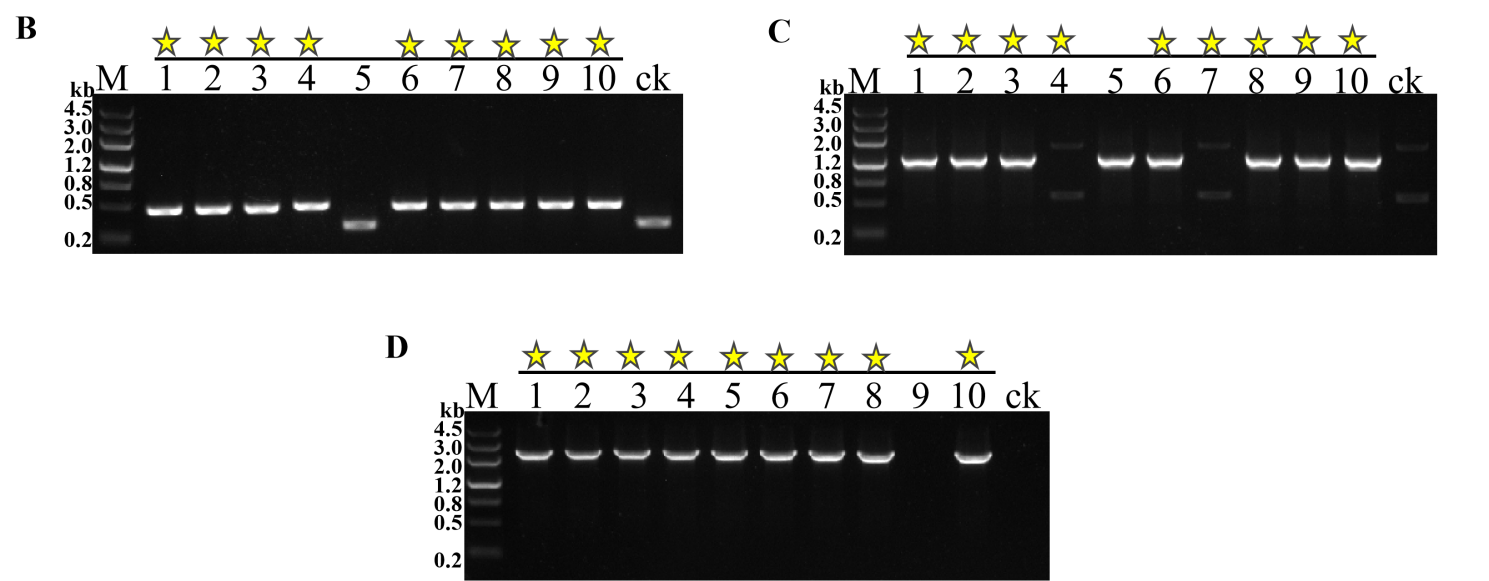

Fig. S4. (A) Principle of gene editing by the Bsu-SRP system based on the "Rock-Paper-Scissors" strategy. (B) Construction results of the pBsu-Scissors-*spo0A* plasmid; (C) Construction results of the pBsu-Rock-*amyE* plasmid; (D) Construction results of the pBsu-Paper-*sacB* plasmid. The yellow asterisks represent successful construction.

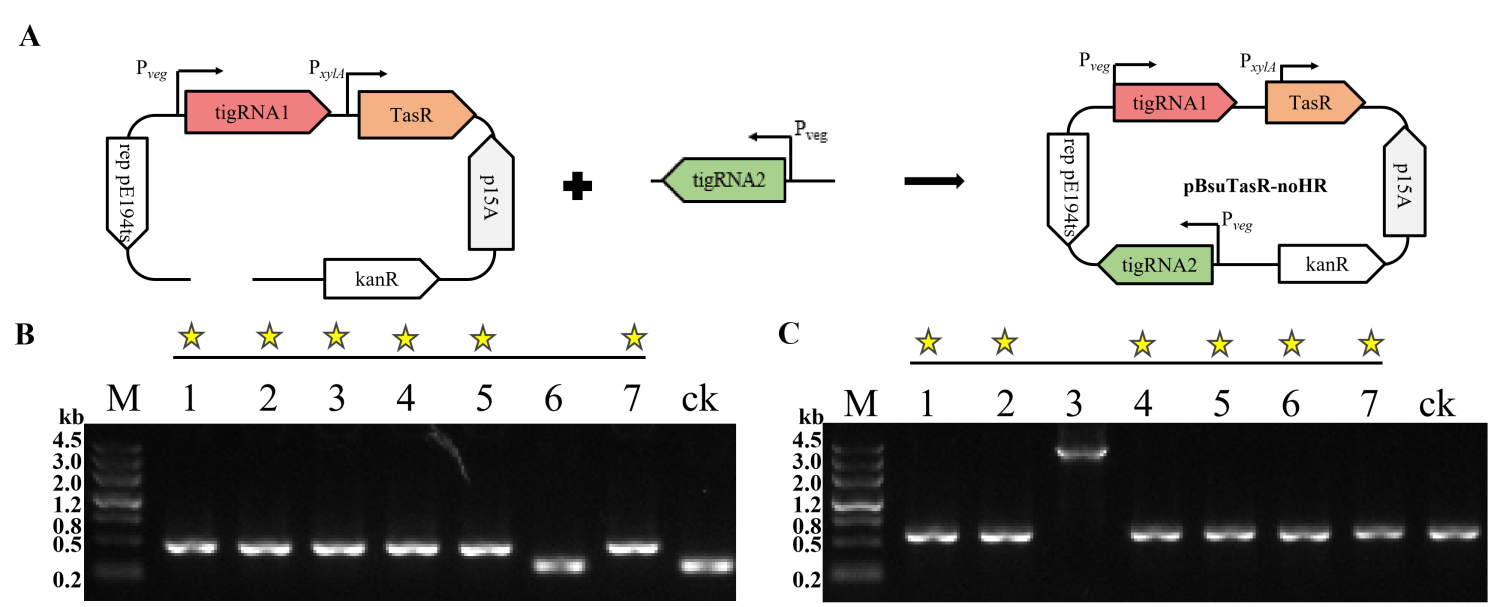

Fig. S5. (A) Workflow for the construction of the pBsuTasR-amyE-tigRNA2 plasmid; (B) Gel image of PCR amplification validating the tigRNA2 portion during construction of the pBsuTasR-amyE-tigRNA2 plasmid; (C) Gel image of PCR amplification validating the removal of the donor DNA portion during construction of the pBsuTasR-amyE-tigRNA2 plasmid. The yellow asterisks represent successful construction.

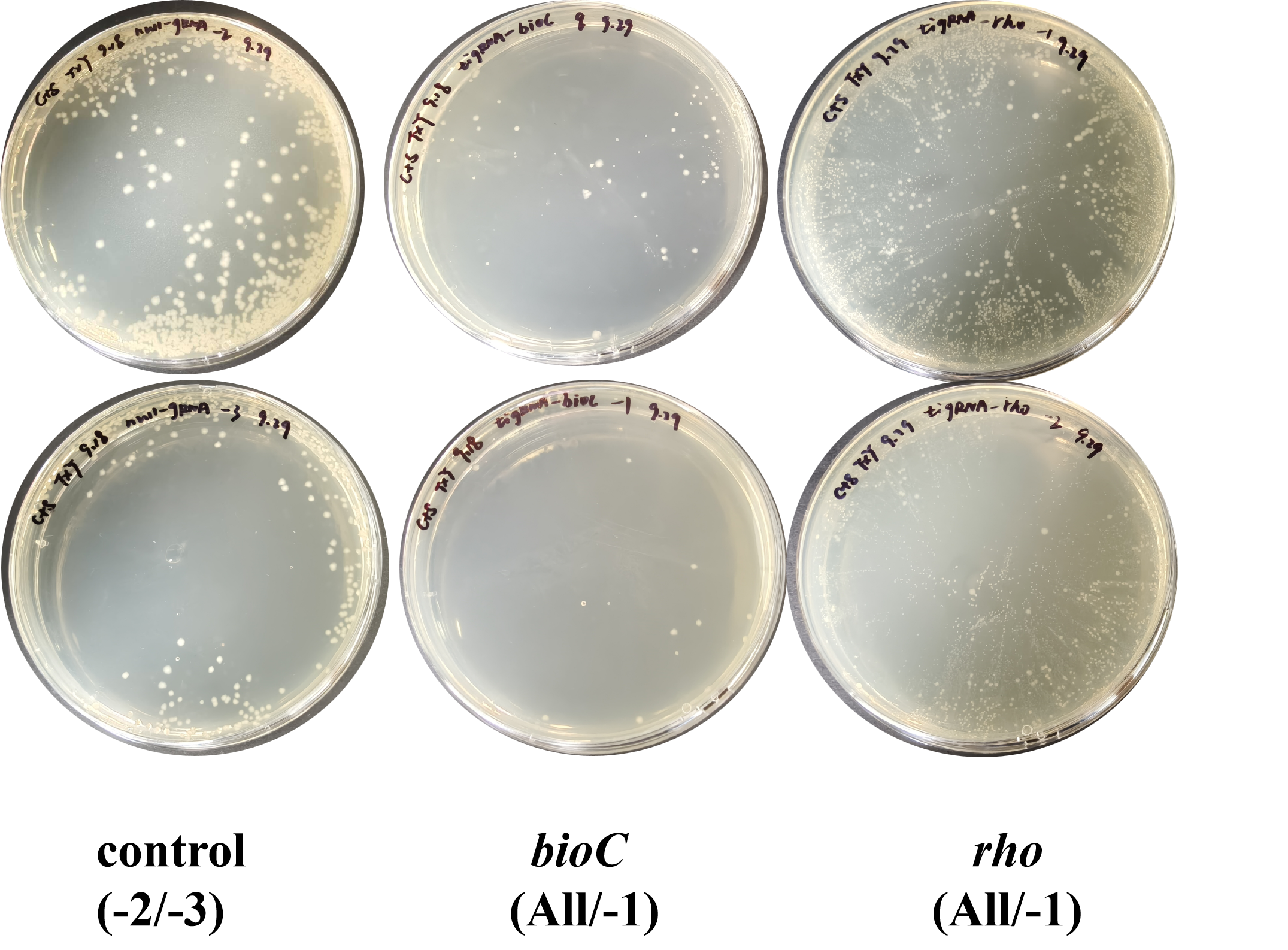

Fig. S6. The testings of cleavage activity of TasR in *E. coli* MG1655 by using pEc-TasR.
